# Quantitative study of the somitogenetic wavefront in zebrafish

**DOI:** 10.1101/419705

**Authors:** Weiting Zhang, Bertrand Ducos, Marine Delagrange, Sophie Vriz, David Bensimon

## Abstract

A quantitative description of the molecular networks that sustain morphogenesis is one of the challenges of developmental biology. Specifically, a molecular understanding of the segmentation of the antero-posterior axis in vertebrates has yet to be achieved. This process known as somitogenesis is believed to result from the interactions between a genetic oscillator and a posterior-moving determination wavefront. Here we quantitatively study and perturb the network in zebrafish that sustains this wavefront and compare our observations to a model whereby the wavefront is due to a switch between stable states resulting from reciprocal negative feedbacks of Retinoic Acid (RA) on the activation of ERK and of ERK on RA synthesis. This model quantitatively accounts for the near linear shortening of the post-somitic mesoderm (PSM) in response to the observed exponential decrease during somitogenesis of the mRNA concentration of a morphogen (Fgf8). It also accounts for the observed dynamics of the PSM when the molecular components of the network are perturbed. The generality of our model and its robustness allows for its test in other model organisms.

## INTRODUCTION

Somites are bilaterally paired epithelial segments of mesoderm that form along the head-to-tail axis of the developing vertebrate embryo. Beginning caudally to the otic vesicle they form periodically from the presomitic mesoderm (PSM) at a time interval that is species specific: about 25 minutes in zebrafish embryos, 90 minutes in chicken, 4-5 hours in humans. Similarly, the number of somite pairs differs among various species. There are 31 pairs of somites in zebrafish, 50 in chicken and several hundred in snakes. As precursors of the vertebrae and skeletal muscles, somites are transient structures: their abnormal formation leads to skeletal and muscular malformations.

Somitogenesis is the process of segmentation of the antero-posterior axis. In zebrafish this process starts at about 10 hpf and ends at 24 hpf. During that developmental interval, as the embryo elongates pairs of somites synchronously and periodically pinch off from the anterior part of the PSM in an anterior to posterior series until 31 pairs of somites are formed. The period between somites, but surprisingly not their size is strongly temperature dependent.

The formation of somites in zebrafish can be divided into two main phases. The first phase is the establishment of segmental pre-pattern in the anterior PSM accomplished by a stripe of gene expression that is believed to be the result of the interactions between a genetic oscillator and a posterior-moving determination wavefront. This pre-pattern determines the place of the morphological segment of a somite which will be formed during the next phase. The Clock and Wavefront model first proposed by Cooke and Zeeman in 1976 (Cooke and Zeeman, 1976), is currently used to describe the output of the complex genetic network underlying the pre-pattern of somite formation. This model proposes that periodic genetic oscillations (the segmentation clock) that move anteriorly (in the PSM reference frame (Dequéant and Pourquié, 2008)), pass through a determination wavefront moving posteriorly (in both the lab and the PSM reference frames) and stop oscillating. As a result a stripe of genes such as Mesp2 are activated to establish the future boundary of the following somite. In this model, the size of a somite is determined by the distance travelled by the determination wavefront during one cycle of the segmentation clock (Dequéant and Pourquié, 2008), see Fig.1.

**Fig. 1:**
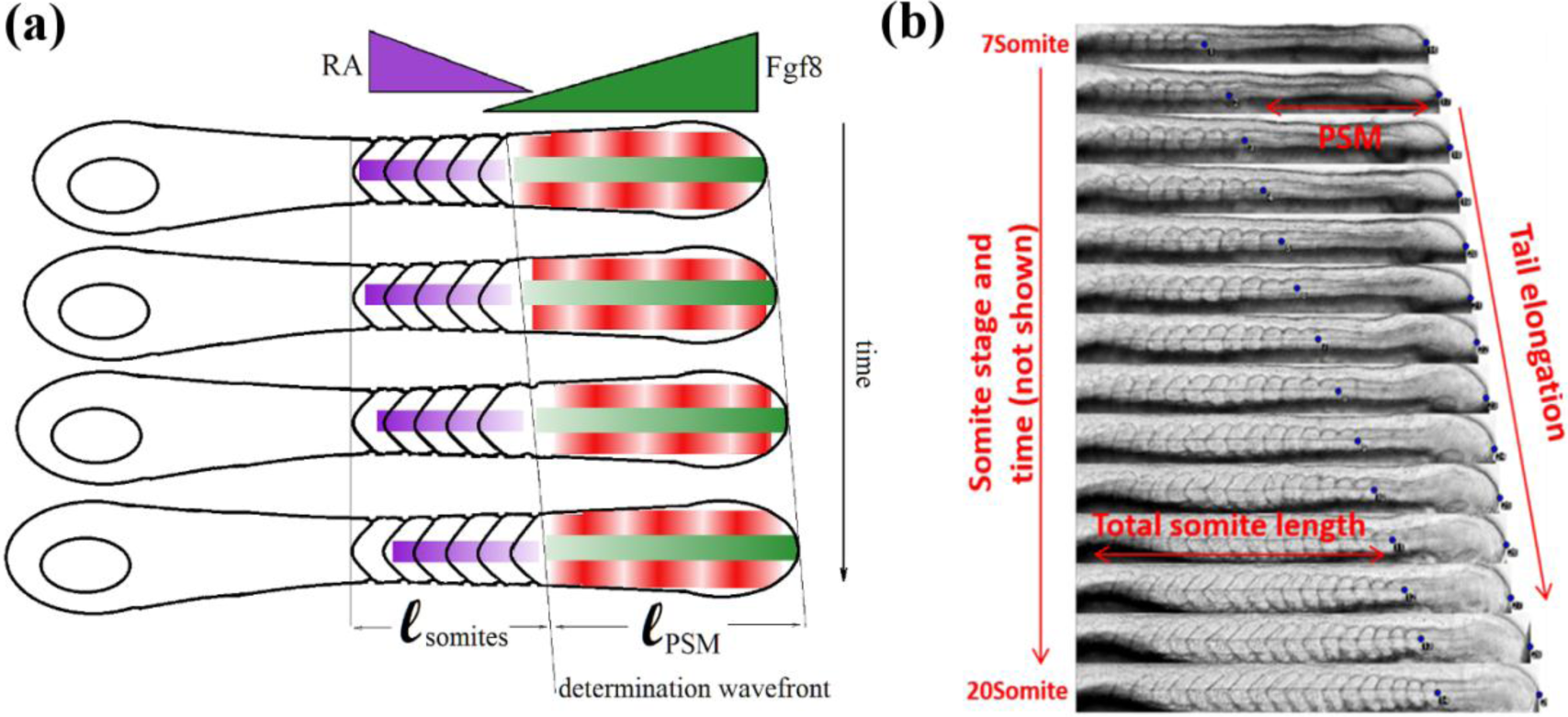
(a) A Clock and Wavefront model: antagonistic gradient of Fgf8 (originating from the posterior PSM, green) and RA (originating from the somites, violet) define a wavefront which interacts with a particular phase of the segmentation clock (in the PSM, red) to generate somites at periodic times and positions. (b) Kymograph of somitogenesis from 7 to 20 somites. The tail elongates at a constant rate V_tail_ while the PSM shrinks at a roughly constant rate V_PSM_ resulting in a somite wavefront propagating at a rate V_front_= V_tail_-V_PSM_.

The second phase of somitogenesis is the formation of the morphological boundaries and the epithelial somites that differentiate from the anterior PSM (Kulesa and Fraser, 2002). Cells in the posterior PSM are loose mesenchyme. As they move to the anterior PSM their motility slows down (Delfini et al., 2005). The cells positioned anteriorly to the determination front undergo a mesenchymal-epithelial transition and progressively become epithelialized spheres. Such boundary formation is based on the action of adhesion molecules and is the result of complex cellular rearrangements. Various members of the Eph/ephrin family of transmembrane signalling molecules are expressed segmentally in the posterior PSM during somitogenesis. This pattern establishes a receptor/ligand interface at each site of somite furrow formation. The Eph/ephrin system mainly controls the inter-somitic furrow formation (Durbin et al. 1998). Fibronectins and cadherins help to correctly localize the cells along the AP axis (Pourquié, 2001).

The segmentation clock driving the differentiation of the PSM into somites at the determination wavefront, has been amply studied and described (see Dequéant and Pourquié, 2008; Soroldoni et al., 2014). It is not the subject of this work. Rather we have studied the molecular network behind the determination wavefront. In contrast with the segmentation clock whose details are species dependent, the main actors of the wavefront (e.g. Fibroblast Growth Factors (FGF), Retinoic Acid (RA)) are conserved in vertebrates (from fish to mammals, including snakes and amphibians, see Gomez et al., 2008). Thus, previous studies have shown that a Fgf8 mRNA gradient forms along the PSM peaking at the posterior part (Dubrulle et al., 2001; Sawada et al., 2001; Delfini et al., 2005; Aulehla and Pourquié, 2010). As cells exit from the PSM, they stop transcribing FGF genes. Thus FGF mRNA progressively decays as cells move towards the anterior of the PSM and an FGF gradient is thus formed (Dubrulle and Pourquié, 2004). This FGF mRNA gradient is subsequently translated into a protein gradient and into a MapK activity gradient along the PSM (as ERK, a MapK protein is activated downstream from the FGF receptor) (Sawada et al., 2001; Delfini et al., 2005).

Retinoic Acid (RA) is also implicated in positioning the determination wavefront (Diez del Corral et al., 2003; Moreno and Kintner, 2004). Opposed to the FGF gradient, RA signaling displays an anterior-to-posterior gradient in the anterior-part of the PSM. RA was found in the segmented somites and the anterior-most part of PSM. It is absent in the tail bud and the posterior PSM (Shimozono et al., 2013). RA is synthesized in two sequential oxidation steps from Vitamin A (retinal). The first one, catalyzed by retinal dehydrogenases, converts retinal to retinaldehyde, which is then converted into RA by the catalytic action of retinaldehyde dehydrogenases such as RALDH2 (Niederreither et al., 2003). RA can be degraded and inactivated by oxidation to 4-hydroxy-RA which is catalyzed by the cytochrome Cyp26 (Kam et al., 2012). During somitogenesis, RALDH2 is expressed in the somites and the anterior-most part of PSM while Cyp26 is expressed in the posterior PSM and tail bud (Blentic et al., 2003; Sakai et al., 2001) and is up-regulated by FGF signaling (Pownall and Isaacs, 2010). An anterior to posterior gradient of RA is thus established in the PSM.

Based on these data and others, Goldbeter, Gonze and Pourquié (Goldbeter et al., 2007; Dequéant and Pourquié, 2008) proposed a model (the G_2_P model) whereby the mutual inhibition of RA and FGF would result in a bistability of the FGF pathway. The differentiation of the PSM into somites would then be triggered by the clock-induced transition of cells from a high FGF activated state into a low FGF activated one. In their model RA acts to repress the translation of the Fgf8 mRNA gradient into protein (or equivalently induce the degradation of the protein), while Fgf8 activates the degradation of RA via the activation of Cyp26, see Fig.2(a). As a result of this mutual inhibition, the concentration of Fgf8 protein (or equivalently the activity of the MapK (ERK) downstream of its receptor) displays a bistable behavior: for a certain range of Fgf8 mRNA concentrations (i.e. in a certain positional window) the MapK activity can be either high or low, see Fig.2(c). Differentiation into somites is then supposed to be triggered (at a certain phase of the segmentation clock) by the inactivation of MapK in that positional window, i.e. by a shift of the MapK activity domain from a rostral position (the limit of the high activity domain) to a more caudal position (the limit of the low activity domain) as observed in recent experiments (Akiyama et al., 2014).

**Fig. 2:**
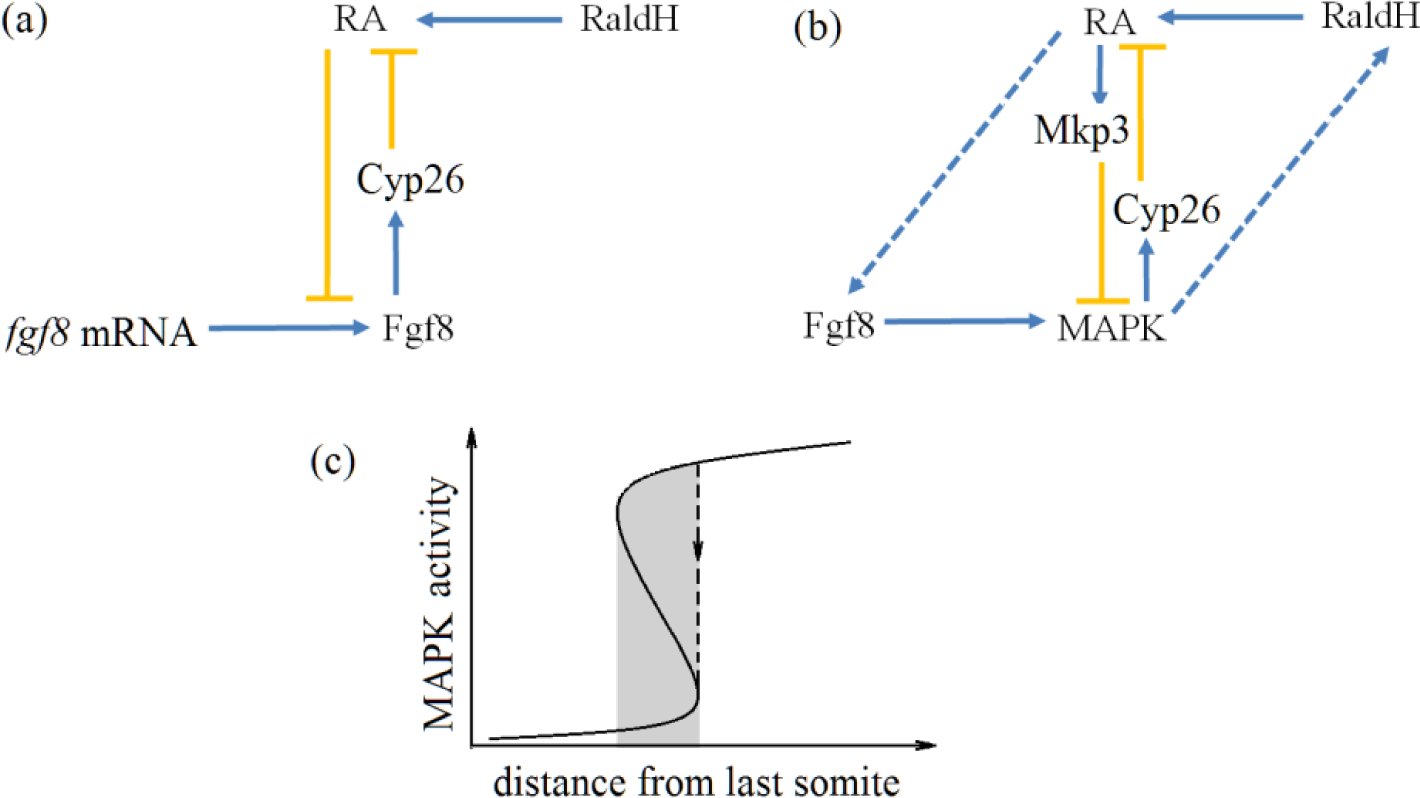
Molecular models of the determination wavefront. (a) The Goldbeter, Gonze and Pourquié (G_2_P) model assumes that RA directly affects the translation of Fgf8 mRNA into protein, while Fgf8 represses RA via the activation of its degradation enzyme Cyp26. (b) The modified version of the G_2_P model (mG_2_P) proposed here takes into account the observed positive feedbacks of RA on Fgf8 and of Fgf8 on RaldH and the mutual inhibition of RA on MapK (via the RA-mediated activation of Mkp3) and of MapK on RA (via the MapK-controlled activation of Cyp26). (c) Both models predict a bistability of MapK activity for a certain positional range (in grey).

However the details of the G_2_P model are contradicted by our own (Fig.S1) and published data (Hamade et al., 2006): RA positively controls Fgf8 and similarly Fgf8 positively controls RaldH (Shimozono et al., 2013). Nonetheless the basic feature of the G_2_P model, namely the bistability of the FGF pathway can be ensured by a negative feedbacks of RA on ERK (mediated by Mkp3 (Dusp6), Moreno and Kintner, 2004) and of ERK on RA (mediated by Cyp26; (Moreno and Kintner, 2004; Pownall and Isaacs, 2010). To take into account these feedbacks we have modified the G_2_P model, see Fig.2(b), and have studied this modified (mG_2_P) model. Details are provided in the Supplementary Material.

In the following we compare the predictions of this mG_2_P model with time-lapse observations of somitogenesis from which we deduce the velocity of the determination wavefront in the moving frame of the PSM. While in the original G_2_P model this velocity is zero, it is in fact negative (the PSM shrinks), probably a result of the decrease with time of the concentration of Fgf8 mRNA during somitogenesis. Measuring the rate of Fgf8 decrease and comparing the rate of shrinkage of the PSM to the predictions of the model provides for a non-trivial test of the model: only an exponential decrease of Fgf8 with time yields the observed almost linear shrinkage of the PSM with time. The model is further tested by comparing its predictions to experiments using various drugs to affect the concentration of RA and impair the activity of Fgf8 and Mkp3 and using conditional expression of an exogenous source of Fgf8 (in an appropriate transgenic line) to increase the Fgf8 concentration. The rate of shrinkage of the PSM, the tail growth rate, wavefront velocity and domain of activity of MapK are measured and compared when possible with the predictions of the model. We find that overall there is a good semi-quantitative agreement between the observations and the predictions of the model.

## RESULTS

### An updated model for the somitogenetic wavefront

The great value of the G_2_P model is to propose a dynamical molecular model for the interaction network between the main known morphogens implicated in somitogenesis: Fgf8 and Retinoic Acid (RA). This allows for a test of the model by a perturbation of these morphogens and their effectors or targets. For example the model assumes a negative feedback of Fgf8 on RA and vice-versa. However we observed that in contradiction with this assumption, RA correlates positively with the expression of Fgf8 (see Fig.S1 and Hamada et al., 2006). Similarly it was shown (Moreno and Kintner, 2004; Hamada et al., 2006) that Fgf8 feeds back positively on the expression of RaldH (the enzyme controlling the synthesis of RA). Hence these details of the G_2_P model are not supported by the experimental data.

Here we propose that the reciprocal negative feedbacks required by the G_2_P model in order to explain the bistability window of the somitogenetic wavefront are implemented at the MapK activity level. Erk, a MapK, is activated (through phosphorylation, Delfini et al., 2005) upon binding of Fgf8 to its receptor (FgfR). Erk is also known to be dephosphorylated (inactivated) by Mkp3 (Dusp6) which has been reported to be under positive control by RA (Moreno and Kintner, 2004). Similarly, Erk has been shown to be implicated in the inhibition of RA (Delfini et al., 2005) via its positive control on Cyp26, the RA degrading enzyme (Moreno and Kintner, 2004). Hence the bistability of the Fgf8 pathway is ensured by the mutual inhibition of RA on Erk and vice-versa. A dynamical molecular model implementing these known observations (hereafter referred to as the modified G_2_P (or mG_2_P) model) is shown in Fig.2(b) and detailed in the Supplementary Information. Like the G_2_P model, it displays a window of bistability along antagonistic gradients of RaldH and Fgf8 mRNA. The presence of a bistability window is very robust to changes in the parameters of the model (some of which can be affected by specific drugs) and to external perturbations of RA and Fgf8.

In the following we will present a detailed quantitative investigation of the somitogenetic wavefront in zebrafish and confront these observations with results of numerical simulations of the mG_2_P model.

### The segmentation clock is insensitive to perturbations of the wavefront

We have studied somitogenesis by time-lapse microscopy (Herrgen et al., 2009) at fixed temperature (T= 26°C) but under a variety of external conditions: with or without specific drugs (DEAB an inhibitor of RaldH, BCI an inhibitor of Mkp3) and with or without perturbations of the morphogens (Fgf8 morpholinos, external RA or induction of exogenous Fgf8). Surprisingly, the time lapse between appearance of somites is unaffected by these diverse perturbations, see Fig.3, which implies that the various actors of the somitogenetic wavefront do not feedback on the clock. This is all the more remarkable considering the extreme dependence of the clock on temperature (Schröter et al., 2008) and the sensitivity of the tail growth rate and PSM shrinkage rate to these perturbations (see below). It would be very interesting to test for the insensitivity of the segmentation clock to these perturbations in other vertebrates (chicken, mouse, etc.).

**Fig. 3.:**
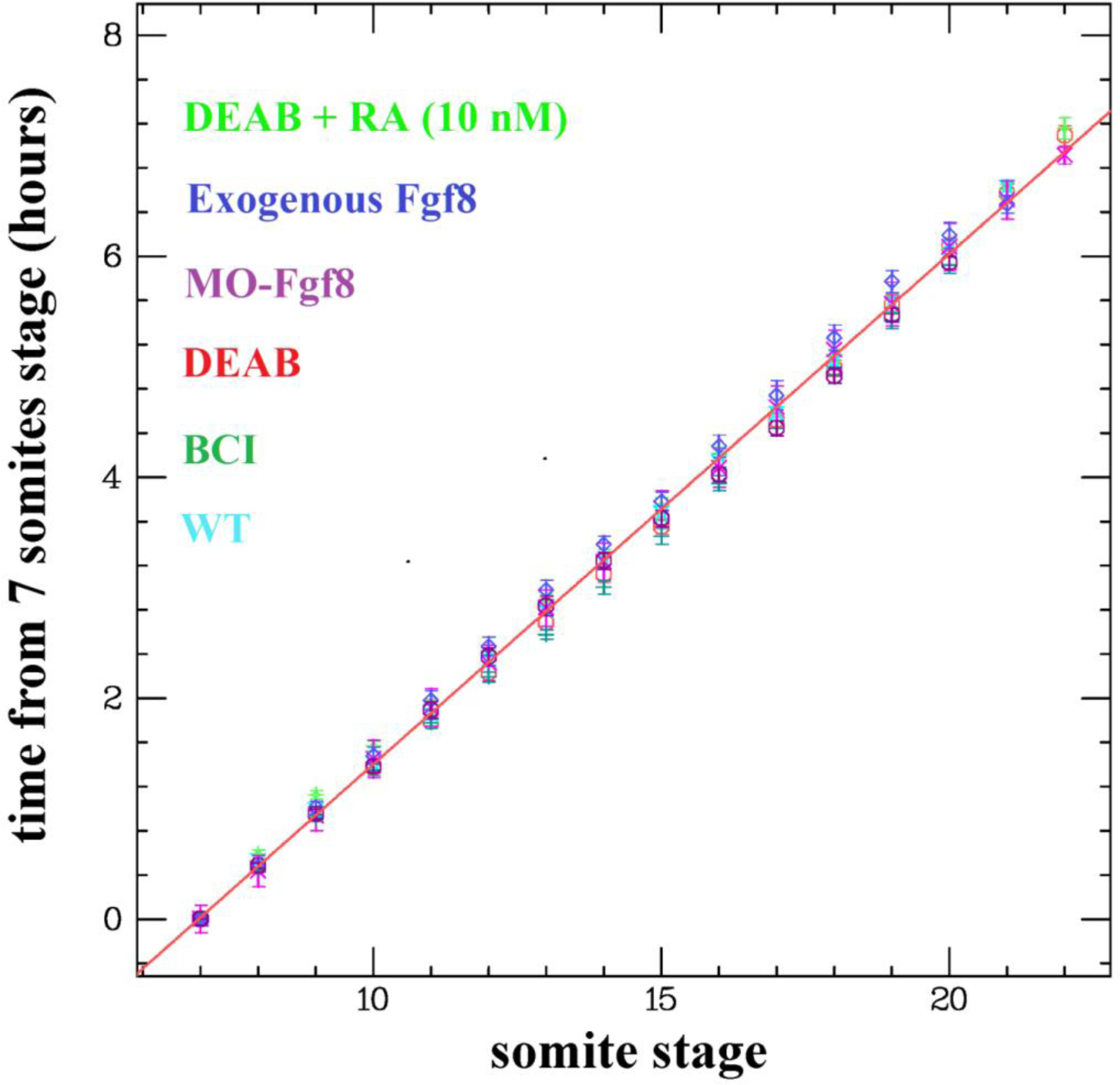
Time of appearance of somites (t=0 at 7 somites stage). Notice the linearity of the plot, i.e. the regularity of the period of somitogenesis, in all the conditions studied here: WT (n=8), DEAB (an inhibitor of RaldH; with (n=14) or without (n=12) external RA), morpholinos against Fgf8 (MO-Fgf8; n=16), BCI (an inhibitor of Mkp3; n=16) or activation of exogenous Fgf8 (n=17).

### The PSM shrinks at a roughly constant rate

Next, we examined the propagation of the wavefront. In the G_2_P model, which is formulated in the tail moving frame, the velocity of the wavefront is equal to the tail growth rate: the position of the bistability window (i.e. the PSM size) is fixed. However, it is a well known fact that the size of the PSM decreases during somitogenesis (Bajard et al., 2014). In the framework of the mG_2_P (and G_2_P) model this can be accounted for by a decrease with time of the Fgf8 gradient during somitogenesis evidenced by a decreasing domain of MapK activity (compare Fig.S2 and S6 (WT) and see Fig.8 below). However the predictions of the models are sensitive to the form of this time decay of the Fgf8 gradient, see Fig.S8. To account for the observed almost linear decrease of the PSM with time (see dots Fig.4(b)), the Fgf8 mRNA gradient has to decay exponentially with time (a linear decay leads to an accelerating shrinking rate). Indeed, qPCR measurements of the Fgf8 mRNA concentration (relative to that of house-keeping genes such as RPL13 or β-actine) suggest that this concentration decays exponentially over the time interval from 5 to 20 somites with a timescale of about 8 somites (8 somitic periods), see Fig.4(a). Rescaling the spatial extent of simulation data by the size of the PSM at 7 somites (t=0) and the timescales by the measured decay of Fgf8 mRNA concentration yields a good agreement between the model and the observed data, see Fig.4(b).

**Fig. 4.**
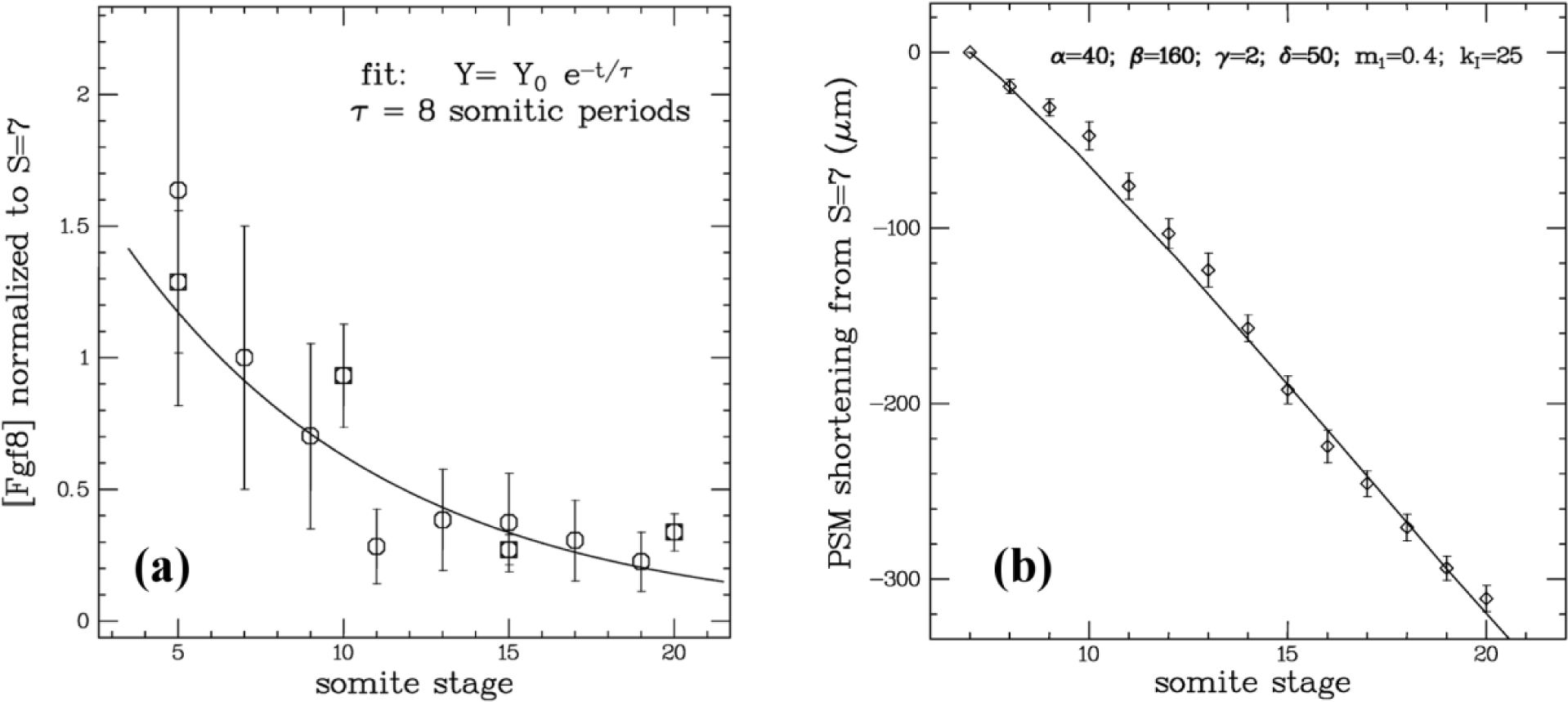
(a) Variation with time of Fgf8 concentration versus somite stage and fit to an exponential decay past 5 somites stage. (b) PSM shortening from 7 somites stage (dots and error bars on mean; n=8) and results (continuous line) of a simulation of the mG_2_P model (with the displayed parameters and assuming an exponential decay of Fgf8 with the timescale measured in (a)). Details in Supp.Mat.

**Fig. 8.**
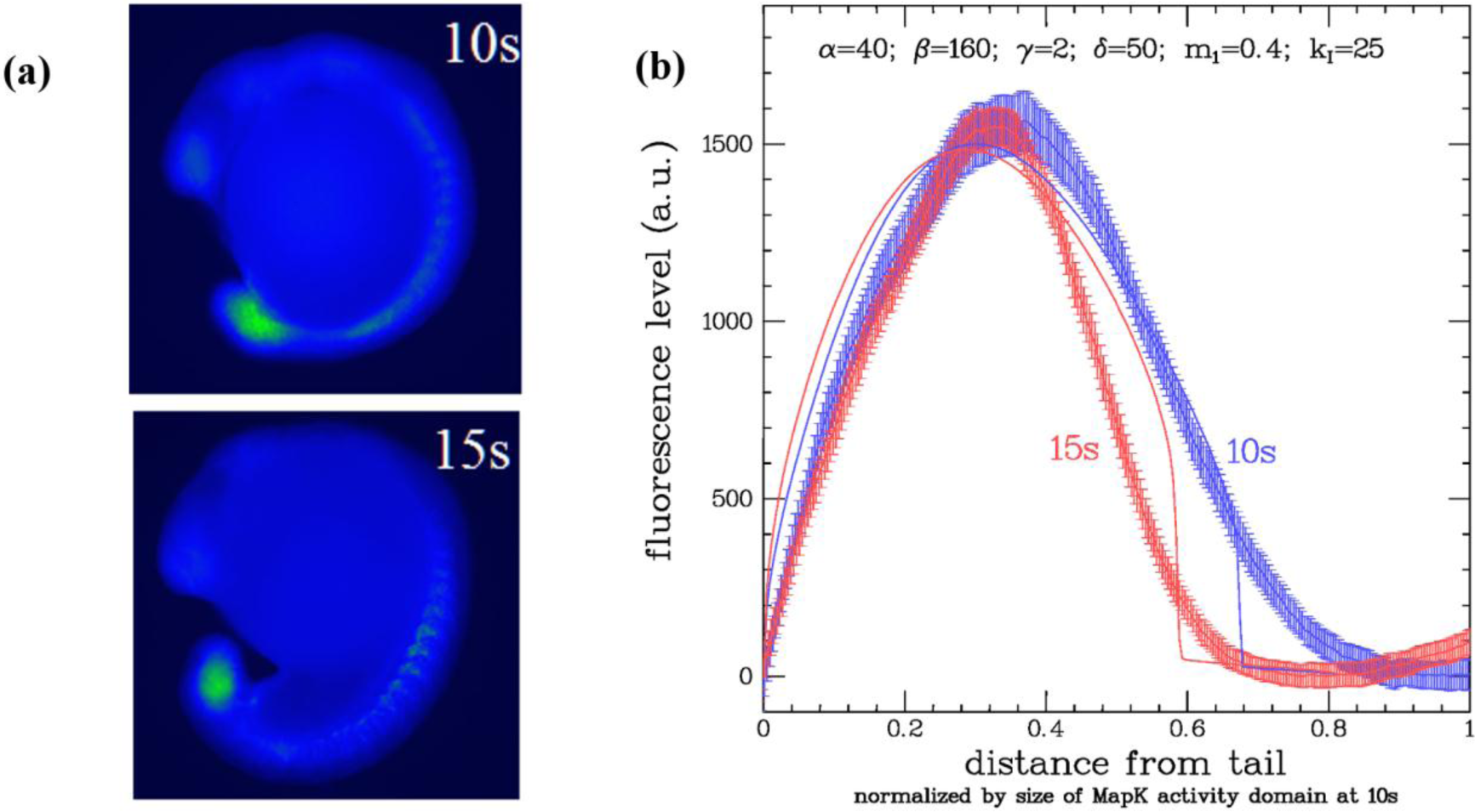
(a) Staining of phosphorylated MapK by antibodies against this active form in WT embryos at 10 and 15 somites. Notice the smaller domain of activity at 15s as compared to 10s. (b) The data in Fig.8(a) was quantified by measuring the fluorescence intensity in a single embryo along the antero-posterior axis and averaged over n=17 (10s) and n=21 (15s) embryos. The averaged data is compared to simulations of the mG_2_P model with the parameters of Fig.4(b) assuming an exponential decrease (see Fig.4(a)) between 7s and 10s (or 15s) of the mRNA Fgf8. The x and y-scales were chosen to fit the data at 10s. The simulation results are in qualitative agreement with the data, even though the latter might not be a perfect reflection of the MapK activity level (which depends on the efficiencies of staining and washing).

The advantage of dynamical molecular models over generic ones (e.g. Turing like models) is that the various actors (molecules, enzymes, etc.) can be acted upon using drugs, RNA interference, genetic knock-in or knock-out methods, etc.. This allows for a qualitative (and sometimes quantitative) test of the models. In this framework we have tested the predictions of the mG_2_P model by acting on many of the actors involved.

### Inhibiting Mkp3 slows down the PSM shrinkage rate and wavefront velocity

First, we looked at the role of Mkp3 (Dusp6). In the mG_2_P model this phosphatase mediates the strength of the negative feedback of RA on ERK (the MapK at the core of the network, see Fig.2(b)) through the dephosphorylation of its active state. We reasoned that inhibiting Mkp3 should reduce the front velocity and more specifically the rate of shrinkage of the PSM, through a decrease in the RA feedback on ERK (dimensionless parameter *k*_*I*_ in the model, details in Supp.Mat.). As can be seen from Fig.5, that is indeed the case: the rate of shrinkage of the PSM decreases in presence of BCI (an inhibitor of Mkp3; (Molina et al., 2009)), in agreement with the prediction of the mG_2_P model. Notice that the presence of the inhibitor also decreases the tail growth rate, so that the overall front velocity is markedly slower in presence of BCI than in its absence (and the resulting somites shorter).

**Fig. 5:**
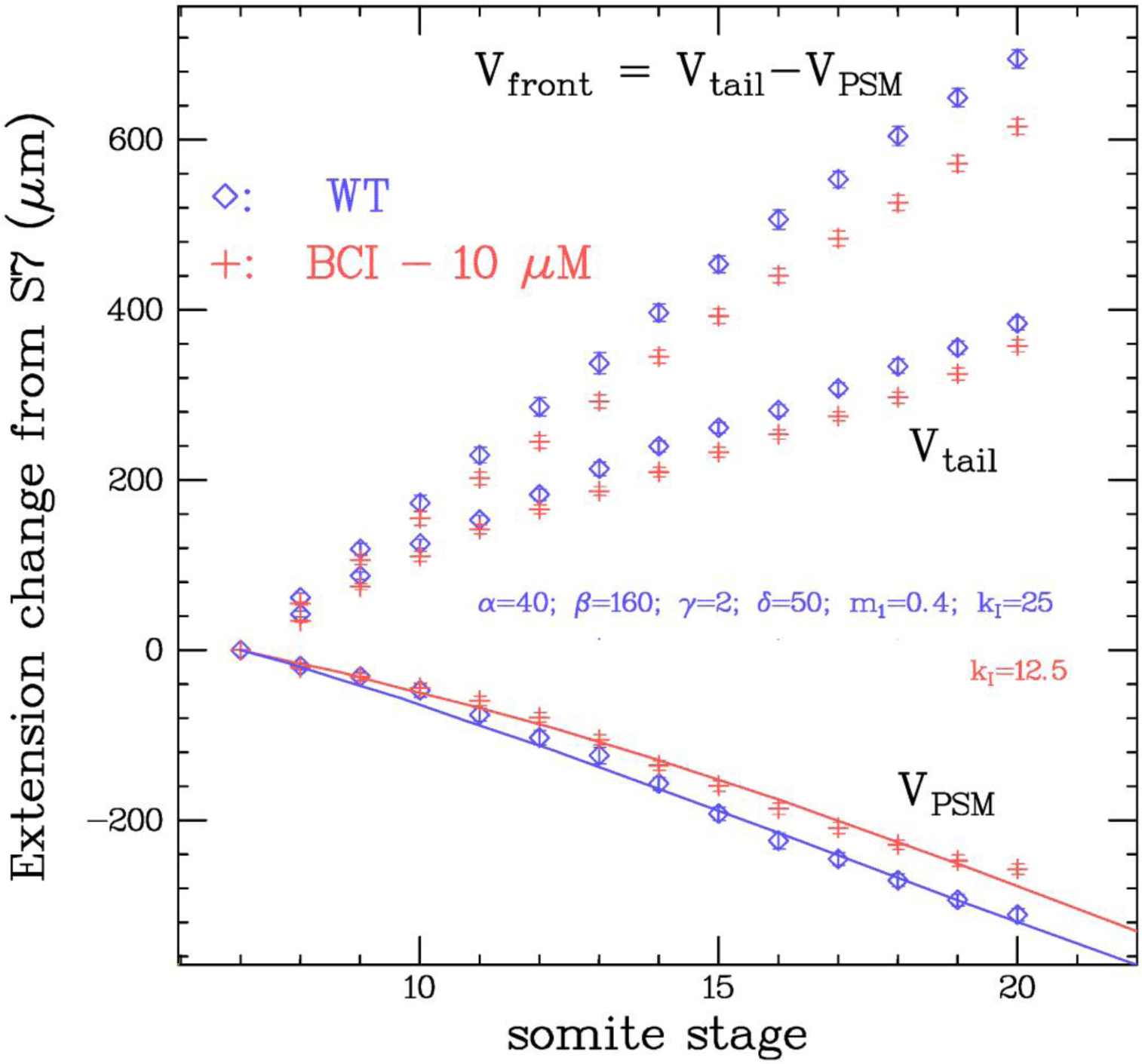
The rates of shrinkage of the PSM (V_PSM_) and growth of the tail (V_tail_) determine the overall somitogenetic wavefront velocity (V_front_=V_tail_ - V_PSM_). In presence of an inhibitor of Mkp3 (BCI), the rates of shrinkage of the PSM and growth of the tail both decrease leading to an overall decrease of the wavefront velocity (dots and error bars on the mean; n=16). Continuous lines: results of simulations of the model with a decreased negative feedback of Mkp3 on ERK (through a factor 2 decrease in *k*_*I*_, details in Supp.Mat.) agree with these observations.

### Inhibiting RA synthesis increases the PSM shrinkage rate but does not affect the wavefront velocity

Next, we examined the role of Retinoic Acid (RA) by inhibiting its synthesis via a drug DEAB known to affect RaldH activity and by simultaneously incubating the embryos in RA. As expected the presence of a RaldH inhibitor led to an increase in the MapK domain of activity, see Fig.S2. Furthermore, as can be seen from the data in Fig.6, the presence of DEAB (with or without the addition of 10nM RA) leads to an increase in the rate of shrinkage of the PSM while simultaneously slowing down the tail growth rate. Surprisingly these two effects compensate each other so that the overall wavefront velocity in presence of DEAB (with or without additional RA) does not affect significantly the wavefront velocity (and thus the somite size) when compared to the WT behavior. This is in contrast with hindbrain formation where the presence of RA rescues the formation of rhombomeres. When comparing the observed data for the rate of PSM shrinkage with numerical simulations of the model a good agreement is observed when assuming a decrease in RalDH activity by a factor two (with or without the presence of a constant external RA source).

**Fig. 6:**
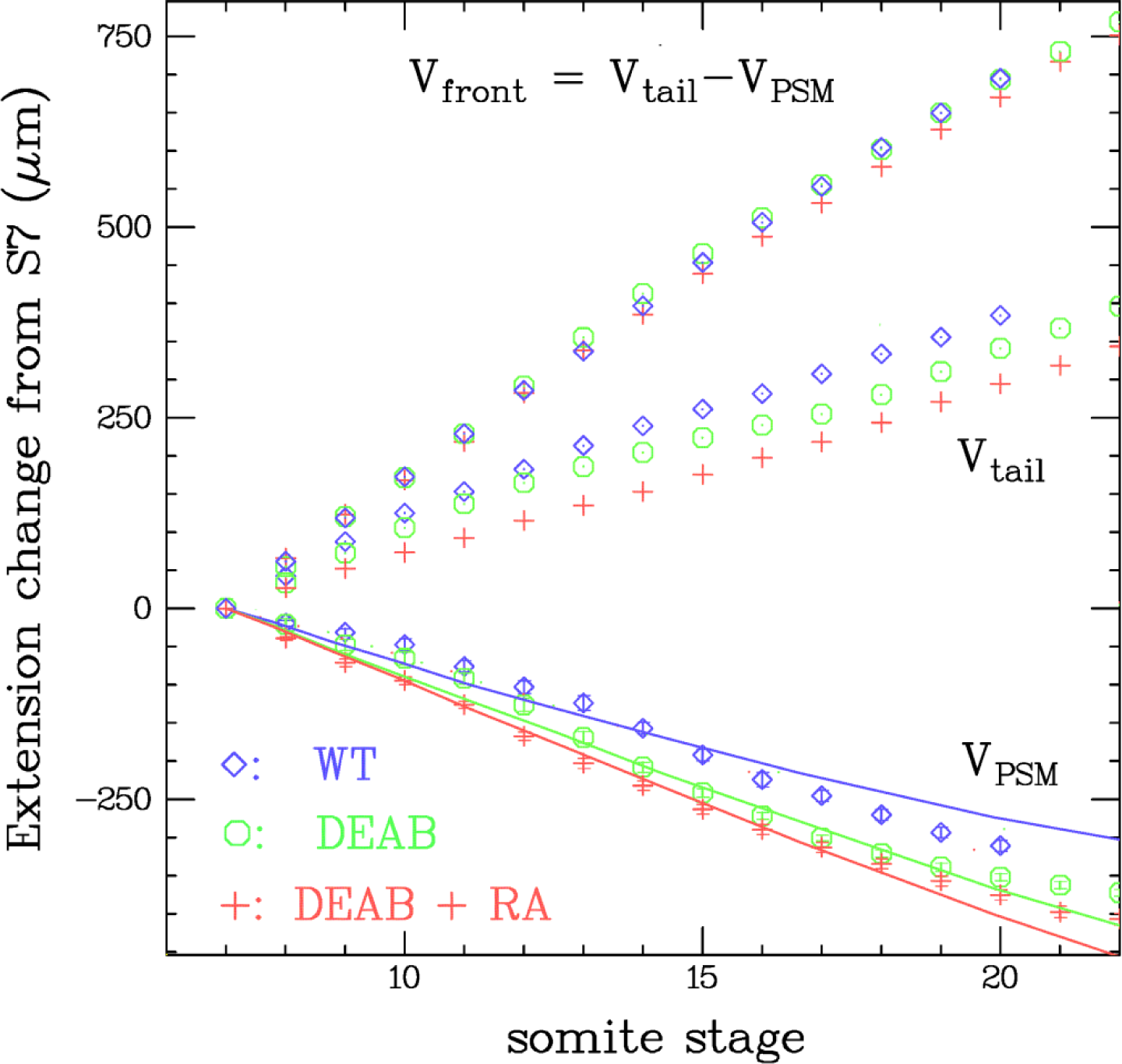
Rates of PSM shrinkage (V_PSM_), tail growth (V_tail_) and wavefront velocity (V_front_) in embryos growing in embryo medium (dots and error bars on mean) without (WT, n=8) or with DEAB (a drug against RaldH) without (n=12) or with addition of 10 nM RA (from 70% epiboly; n=14). Notice however that the wavefront velocity (V_front_ = V_tail_ - V_PSM_) is not affected by these conditions. Continuous lines: results of simulations of the model assuming a decrease of the rate of RA synthesis (parameter α in the model) by a factor two (α = 80 ⟶ 40) in the presence of DEAB and in presence or not of 10nM RA, are in fair agreement with these observations.

### Fgf8 inhibition and overexpression decrease the wavefront velocity but do not affect much the PSM rate of shrinkage

Next, we examined the role of Fgf8 by reducing the concentration of its mRNA (with morpholinos (MO) directed against its mRNA) and by inducing the expression of an exogenous source of Fgf8 at the beginning of somitogenesis. We injected morpholinos against Fgf8 (Draper et al., 2001) at the one cell stage and verified that they affected somitogenesis as previously reported (see Fig.S3). We then compared by time lapse microscopy the course of somitogenesis in embryos injected or not with morpholinos. The results are shown in Fig.7(a). While the injection of morpholinos did reduce, as expected, the size of the MapK activity domain (see Fig.S4), it did not affect significantly the rate of shrinkage of the PSM. It did however decrease the tail growth rate, thus reducing the wavefront velocity, Fig.7(a). In agreement with these observations, simulation of the mG_2_P model with a 60% decrease in the concentration of Fgf8 mRNA does not yield a significant change in the rate of PSM shrinkage.

**Fig. 7:**
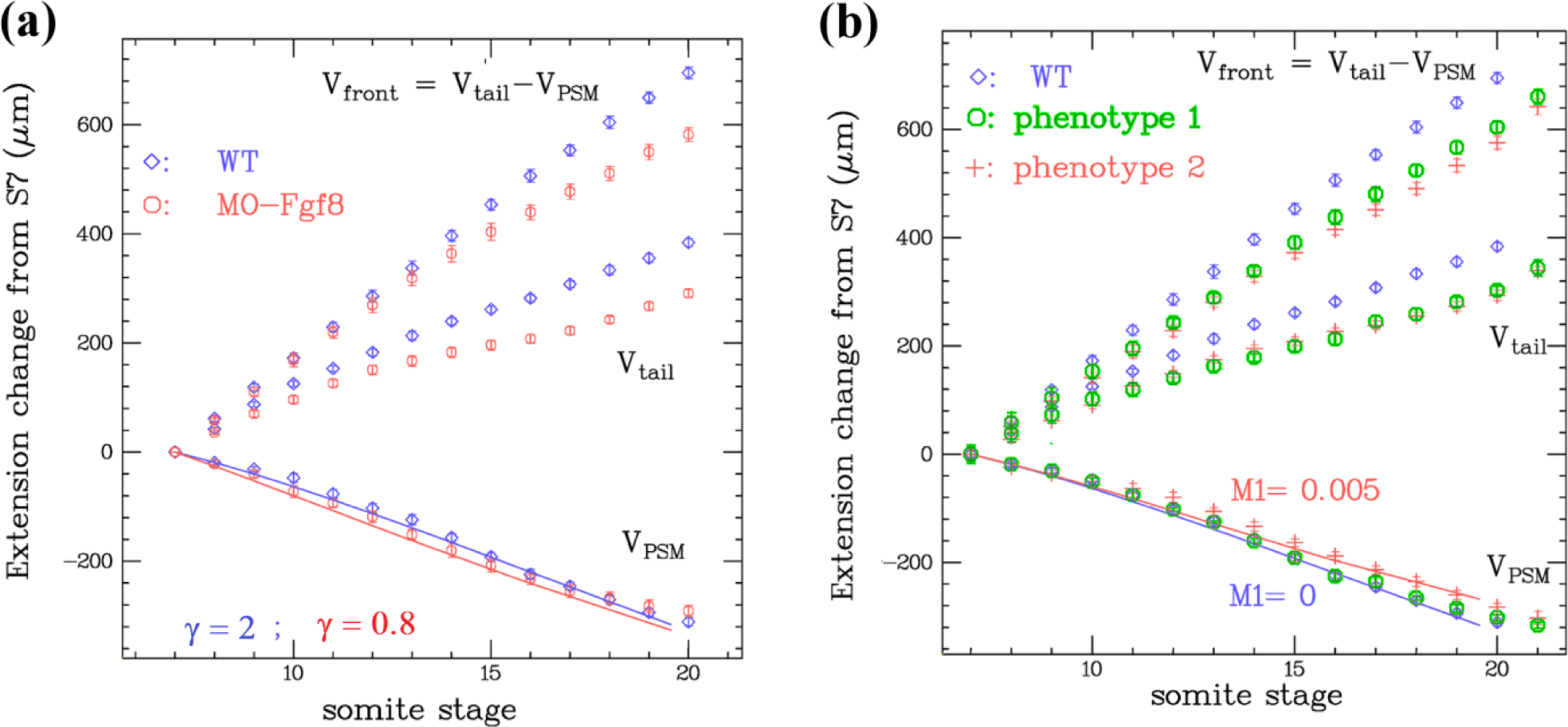
Rates of PSM shrinkage (V_PSM_), tail growth (V_tail_) and wavefront velocity (V_front_) in embryos growing (a) without or with morpholinos against Fgf8 (MO-Fgf8; dots and error bars on mean; n=16) injected at one-cell stage or (b) in which an exogenous source of Fgf8 was turned on (n=17). While V_PSM_ is unaffected by MO-Fgf8 and slightly decreases upon over-expression of Fgf8 (in the strong phenotype 2 embryos), V_tail_ decreases in both conditions, resulting in an overall decrease of V_front_. Continuous lines: simulations of the mG_2_P model with a 60% decrease in Fgf8 mRNA (due to interference with MO-Fgf8; γ=0.8 instead of γ=2 in model, see Supp.Mat.) qualitatively reproduce the data (continuous line in (a)). Similarly simulations with an increasing Fgf8 mRNA of about 2% of the measured increase (see Fig.S7) seem to reproduce the data observed in the strong phenotype 2 case (see Supp.Mat. Fig.S5).

We also examined the effect of the induction of an exogenous source of Fgf8 on somitogenesis. We used embryos that had a Fgf8 gene under control of a UAS promoter that could be induced upon activation of a Gal4-ERT transcription factor. We used incubation in cyclofen (or caged-cyclofen followed by short UV illumination, (Zhang et al, 2018)) to continuously activate this exogenous source of Fgf8. In such conditions we observed a phenotype with a strong deformation of the yolk and smaller somites (see Fig.S5). The MapK domain of activity enlarged (see Fig.S6) as expected from an increase in Fgf8 concentration. Similarly the PSM shrinkage rate was (slightly) reduced, see Fig.7(b), as expected from a continuously increasing Fgf8 mRNA which displaces the instability window rostrally. The expression of exogenous Fgf8 also reduced the tail growth, Fig.7(b), as did the presence of morpholinos against Fgf8, suggesting that the endogenous Fgf8 concentration is optimal for maximal growth of the tail. Results of qPCR show that the induced exogenous source of Fgf8 mRNA increases linearly with time with a doubling of the initial Fgf8 concentration over a 20 somite period (see Fig.S8). Simulations of the mG_2_P model show that the PSM shrinkage rate is reduced (as observed experimentally) and can even be reversed at high enough mRNA production rates. A good fit with the observed data is however obtained with an effective rate of increase of only 2% of the measured rate of increase of exogenous Fgf8 (Fig.S7), which suggests that some mechanisms may limit the translation or export of this exogenous Fgf8.

### The MapK domain of activity in the PSM shrinks during somitogenesis

Finally we compared the measured MapK domain of activity with the one predicted from the numerical simulations of the mG_2_P model. We used Immuno-Histo-Chemistry (IHC) staining with antibodies against the active (phosphorylated) form of ERK to image in fixed embryos the MapK active domain at various stages of somitogenesis. From the images at 10 somites (10s) and 15 somites (15s), see Fig.8(a), it is clear that the MapK domain of activity shrinks with time, as is expected from a decreasing concentration of Fgf8 in the PSM.

## DISCUSSION

We have presented a detailed quantitative study of a model of the somitogenetic wavefront which incorporates many known results under a common theoretical description. The proposed mG_2_P model adopts the concept of bi-stability from the original G_2_P model (Goldbeter et al., 2007). It incorporates elements from Moreno and Kintner (the positive feedback of RA on Mkp3 and Fgf8; see also Hamada et al., 2006 and Shimozono et al., 2013). (the positive feedback of Fgf8 on RaldH) and puts at its core a Mapk (ERK) which switching from phosphorylated to dephosphorylated state has been amply documented as a key step in early differentiation into somite fate (as early as S-IV; (Delfini et al., 2005; Sawada et al., 2001; Akiyama et al., 2014). In the proposed model the bi-stability is a result of the mutual inhibition between RA and MapK. Many of the results presented here on the response of the domain of MapK activity to various perturbations have been qualitatively reported previously (such as the inhibition of the FGF pathway with SU5402 or its activation with implanted Fgf8 soaked beads, see (Delfini et al., 2005; Sawada et al., 2001).

What has not been attempted until now was a quantitative comparison between a detailed molecular model of the wavefront and the dynamics of PSM shortening under various perturbations. We have chosen to conduct that comparison in zebrafish, a model animal that has the advantage of quick development (somitogenesis occurs overnight), ease of transgenics and the possibility to monitor simultaneously the development of many (15-20) embryos in the same conditions (allowing for better statistics). In contrast with other organisms (mouse, chicken, snake; Gomez et al., 2008), in WT zebrafish embryos the PSM shrinks at roughly constant rate between 7 to 25 somites (Gomez et al., 2008, Schröter et al., 2008). We measured that rate in WT embryos and in embryos subjected to a variety of perturbations: inhibitors of Mkp3 and RaldH (with or without addition of external RA), morpholinos against Fgf8 and induction of an exogenous source of Fgf8. In these various conditions we noticed that the rate of shrinkage was still quite constant but differed from the rate in WT conditions.

We compared our observations to simulations of the mG_2_P model in the adiabatic approximation. Namely we attempted to explain the shrinkage of the PSM by assuming that the Fgf8 mRNA at the tail end decayed on a timescale larger than the relaxation time (less than a somitic period) of the network describing the wavefront (Fig.2(b)). Using qPCR to study experimentally the variation with time of the Fgf8 mRNA we observed that it decayed exponentially with a long timescale (about 8 somitic periods, i.e. ∼4h) justifying our assumption. In this adiabatic approximation, the mG_2_P model can be reduced to a dimensionless form that depends on 6 parameters only (instead of 18 for the full dynamics). Variation of some of these parameters with distance from the tail (e.g. gradients of RaldH and Fgf8 mRNA) results in the possibility of having two co-existing stable states (high and low MapK activity) in a certain range of distances from the tail end, that define the position of the wavefront. We have not attempted to find the best fit of our 6 parameters to the observed data but have shown that for some reasonable values of the parameters and using the measured exponential decay of Fgf8 mRNA with time a fair quantitative agreement is obtained between the results of simulations and the observations of the PSM shrinkage under a variety of perturbations.

A sensible critique of this approach is that with 6 parameters the mG_2_P model could be fiddled to fit any data. Fortunately, that is not the case as the dynamics of the PSM shrinkage in this model is quite robust (or insensitive) to variation of the parameters. For example, with a linear decrease with time of the Fgf8 mRNA, we could not find parameters for which the PSM shrinks almost linearly with time, see Fig.S8(b,d). The model can thus be used to falsify various hypotheses.

For example a linear gradient of Fgf8 mRNA only can sustain bistability (see Fig.S8(g)) and could account for the formation of somites in absence of an RA gradient (as reported by Niederreither et al., 2002). However the almost constant rate of shrinkage of the PSM observed in WT embryos is incompatible with a linear gradient of Fgf8 mRNA only. Similarly linear gradients of Fgf8 mRNA and of the rate of Cyp26 transcription while compatible with many of our observations (see Fig.S9) cannot account for the observed faster rate of PSM shrinkage in presence of a RaldH inhibitor (DEAB).

Due to its insensitivity to variation of its parameters, the model can be easily falsified (i.e. it cannot be fiddled to fit any data). For example in chicken and snake the PSM extension varies in a non-monotonous fashion (it increases and then decreases; Gomez et al., 2008). Within the present model this result suggests that the Fgf8 mRNA could increase initially and then decrease. By measuring the time dependence of the various actors of the network (in particular the Fgf8 mRNA concenration), and using that data as an input to a simulation of the mG_2_P model (as we have done here with the observed exponential decay of Fgf8 mRNA) one could compare the simulated PSM dynamics to the observed one and test for the validity of the mG_2_P model in these other organisms.

Finally, it is interesting to notice that similar antagonistic interactions between RA (high in the anterior part of the embryo) and MapK (active in the posterior end) have been reported in an invertebrate chordate where they control posterior patterning (Pasini et al., 2012).. It is tempting to speculate that in vertebrates, segmentation evolved via the inhibition of this primitive network by an independent pulsating one (the segmentation clock). The observed insensitivity of the zebrafish somitogenetic period to perturbations of the actors of the wavefront (a robustness that should be tested in other organisms) is compatible with this hypothesis.

## MATERIALS AND METHODS

### Fish lines and maintenance

Zebrafish were raised and maintained on a 14 h-10 h light-dark diurnal cycle with standard culture methods (Westerfield, 2000). Embryos collected from natural crosses were staged according to Kimmel (Kimmel et al., 1995). The *Tg(uas:fgf8-T2A-cfp;ubi:eos)* was generated by injecting the plasmid *pT24-uas:fgf8-T2A-cfp;ubi:Eos* (a gift from Michel Volovitch) which contains the homologous cDNA sequence of *fgf8* from Danio rerio, with Tol2 mRNA transposase. Founder transgenic fish were identified by global expression of Eos. The *Tg*(*ubi:Gal4-ERT;cry:CFP)* was described in (Feng et al., 2017). The double transgenic line *Tg(ubi:Gal4ERT;cry:CFP; uas:fgf8-T2A-cfp;ubi:Eos)* was created by crossing *Tg(uas:fgf8-T2A-cfp;ubi:eos)* and *Tg*(*ubi:Gal4-ERT;cry:CFP).* Founder double transgenic fish were selected by global expression of Eos and expression of CFP in the developing eyes.

### Morpholino microinjection

A morpholino targeting Fgf8 (MO-Fgf8) was used to alter the endogenous Fgf8 pre-mRNA splicing. With sequence 5’-TAGGATGCTCTTACCA<u>TGAACGTCG</u>-3’ (Draper et al., 2001), it spans the junction of exon3/intron3 of Fgf8 (intronic sequences underlined). It displays 9 mismatches with the exogenous Fgf8 in *uas:fgf8-T2A-cfp* (mismatch sequences underlined 5’-TAGGATGCTCTTACCA<u>TGAACGTCG</u>-3’). Zebrafish embryos were injected at the one to four cells stage with 1, 2,4, 8 nl of the solubilized compounds contained 1ng/ul MO-Fgf8 as final concentration in 1x Danieau buffer (58mM NaCl, 0.7mM KCl, 0.4mM MgSO_4_, 0.6 mM Ca(NO_3_)_2_, 5.0 mM HEPES pH 7.6). Embryos were imaged for phenotypic analysis at 24 hpf and then fixed with PAXgene Tissue Container Product (Qiagen) for RT-PCR.

### RT-PCR

Wild type embryos uninjected or injected with various amount of MO-Fgf8 were fixed at 24 hpf using PAXgene Tissue Container Product (Qiagen). Extraction of total RNA was performed by using RNeasy Micro Kit (Qiagen), according to the manufacturer’s instructions. cDNA synthesis was performed using sensiscript reverse transcriptase (Qiagen) with an anchored Oligo(dT)23VN primer (NEB). PCR was performed using the Phusion High-Fidelity DNA Polymerase (NEB) with the following protocol: 98°C for 30 s; then 30 cycles (98°C for 10 s, 60°C for 30 s, 72°C for 30 s); 72°C for 10 min. A pair of primers, P1 (5’-ACCATTCAGTCCCCGCCTAA-3’) and P3(5’-GCCAATCAGTTTCCCCCTCC-3’) which respectively match exon3 and exon4 of Fgf8, were used to detect the expression of the pre-mRNA of the endogenous Fgf8 resulting in a 1900 bp PCR product and the expression of the exogenous and correctly spliced Fgf8 resulting in a 283 bp PCR product. Another pair of primers, P3 (same as before) and P2 (5’-CCCCTCCGTTTGAACCGTAA-3’) against intron3 of Fgf8, was used to analyze the expression of the pre-mRNA of the endogenous Fgf8 with a PCR product of 722bp. This pair cannot amplify the exogenous and correctly spliced Fgf8. Actb2 was amplified as a reference gene with the following primers: 5’-TGTACCCTGGCATTGCTGAC-3’ and 5’-CCATGCCAATGTTGTCGTTTG-3’.

### Quantitative RT-PCR

Wild type embryos (5-10 embryos in each group) were fixed at the somite stages mentioned in the text. Embryos fixation and total RNA extraction were performed as mentioned above. cDNAs were synthesized using Fluidigm Reverse Transcription Master Mix kit. PCR was run on Light cycle 480 Real-Time PCR System (Roche) with the following protocol: 50°C for 2 min; 95°C for 10 min; 45 cycles (95°C for 15 s, 60°C for 1 min); hold at 37°C. RT-qPCR was performed using the aforementioned cDNAs with TaqMan Universal PCR Master Mix and TaqMan Gene Expression Assay: Dr03119263_m1(rpl13a) and Dr03105657_m1(fgf8) (Applied Biosystems). The difference (ΔCp) in the number of amplification cycles between these genes (in various conditions) and a reference gene (rpl13a) was measured in duplicate PCR assays. The relative gene quantification was calculated by 2^-^ΔΔ^Cp^ = 2^-(^Δ^Cp sample-^Δ^Cp rpl13a)^.

### Immunohistochemistry on zebrafish embryos

The embryos obtained in the various conditions and the different stages mentioned in the text were fixed in 4% PFA overnight at 4°C, followed by dehydration with 100% methanol at −20°C for more than 1 day. After gradual rehydration of methanol and wash with PBS/Tween 0.1%, the embryos were incubated in a blocking solution: 1%Triton, 1% DMSO, 1% BSA and 10% sheep serum (Sigma) in PBS on a shaker for 1 hour at room temperature, followed by incubation with 1:400 Phospho-ERK Monoclonal Antibody as the primary antibody (Thermo, MA5-15173) on a shaker overnight at 4°C. After extensive washing with PBS/Tween and incubation in blocking solution, a second antibody (anti-Rabbit conjugated to Alexa Fluor 488, Invitrogen) diluted 700x was added overnight at 4°C. After washing with PBS/Tween, images were taken on a Nikon Ti microscope equipped with a Hamamatsu Orca camera.

### Whole-mount In situ hybridization

Whole-mount in situ hybridization with digoxigenin-labeled riboprobes was performed as described previously (Thisse et al., 1993). The antisense riboprobes were synthesized from template plasmids (gift of P. Rosa, IBENS) containing fgf8 full length cDNA (Fürthauer et al., 1997).

### Drug treatments

Wild type embryos were incubated in 10 μM DEAB (with or without addition at 70% epiboly of 10 nM all-trans RA), or BCI 10 μM (all Sigma-Aldrich; all from a 10 mM stock in DMSO diluted with embryo medium (EM)). The double transgenic embryos *Tg(ubi:Gal4ERT;cry:CFP;uas:fgf8-T2A-cfp;ubi:Eos)* were incubated at 70% epiboly in 3 μM cyclofen (gift of L. Jullien, see Sinha et al., 2010) diluted in EM. As controls, siblings were kept in EM.

### Fluorescent microscopy

All fluorescent images were taken on a Nikon Ti microscope equipped with a Hamamatsu ORCA V2+ camera and a 10x plan fluo objective. Filter setting of CFP: excitation at 438 nm, emission at 483 32 nm; mEosFP and Alexa 488: excitation at 497 nm, emission at 535 nm.

### Time lapse video

All embryos were dechorinated before bud stage using acute tweezers. The embryos were mounted in an agarose mold and kept in a temperature incubator at 26°C during imaging, following published protocols (Herrgen et al., 2009). To prevent evaporation during recording caused by the difference in temperature between the microscope room and the incubator, mineral oil (Sigma) was add to cover the surface of the medium. Time-lapse video was performed on a Nikon Ti Microscope with a 10x plan fluo objective. Images were taken by an ORCA V2+ Camera (Hamamatsu). The entire microscope system including the filter rotor, the motorized stage and the camera was driven by Micro-Manager. Up to 20 embryos per experiment were scanned at an interval of 5 minutes. For each embryo, a Z-stack with 6 planes at 25μm intervals was recorded.

### Image analysis of Time lapse video

Time lapse images were analyzed by FIJI. The Gaussian-based stack focuser in FIJI Time Lapse plugin was used to choose and combine the focused areas from different Z-positions of one time point. The measurement of tail, PSM and somite length was done using the measurement function in FIJI.

## Acknowledgements

We thank A.Oates and D.Soroldini for their help and support with time-lapse microscopy on zebrafish embryos. We further acknowledge C.Gauron and M.Volovitch for their help in the construction of transgenic fish and L.Jullien for his support with cyclofen and its derivatives. We thank them all and A.Goldbeter and V.Hakim for constructive criticism during the conduction of this work. This work was partially supported by grants ANR-10-LABX-54 MEMO LIFE and ANR-11-IDEX-0001-02 PSL* Research University and PSL grants SuperLINE and MicroGUT. The authors declare that they have no competing interests.

## Authors contributions

W.Z. and D.B. planned the experiments. W.Z. performed most of the experiments. D.B. developed the model and simulated it. B.D. and M.L. performed some of the RT-qPCR experiments, S.V. developed some of the zebrafish lines used here.

## SUPPLEMENTARY MATERIAL

**Fig. S1:**
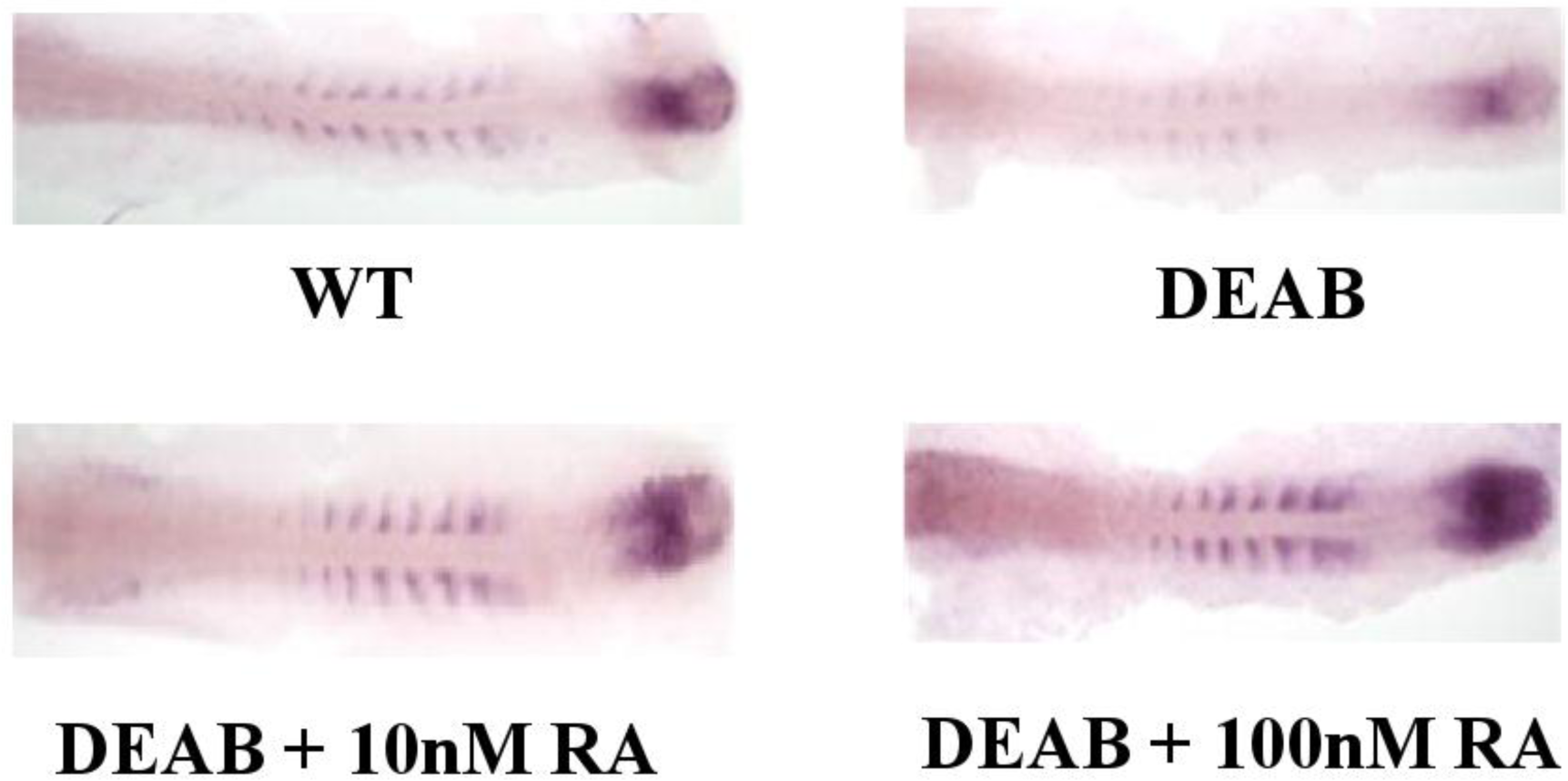
In-situ hybridization against Fgf8 mRNA in WT embryos and in embryos incubated in 10 μM DEAB (from one cell stage) without or with 5 minute incubation at 90% epiboly in 10 or 100 nM Retinoic acid. Notice the positive correlation between the levels of RA and Fgf8 expression.

**Fig. S2:**
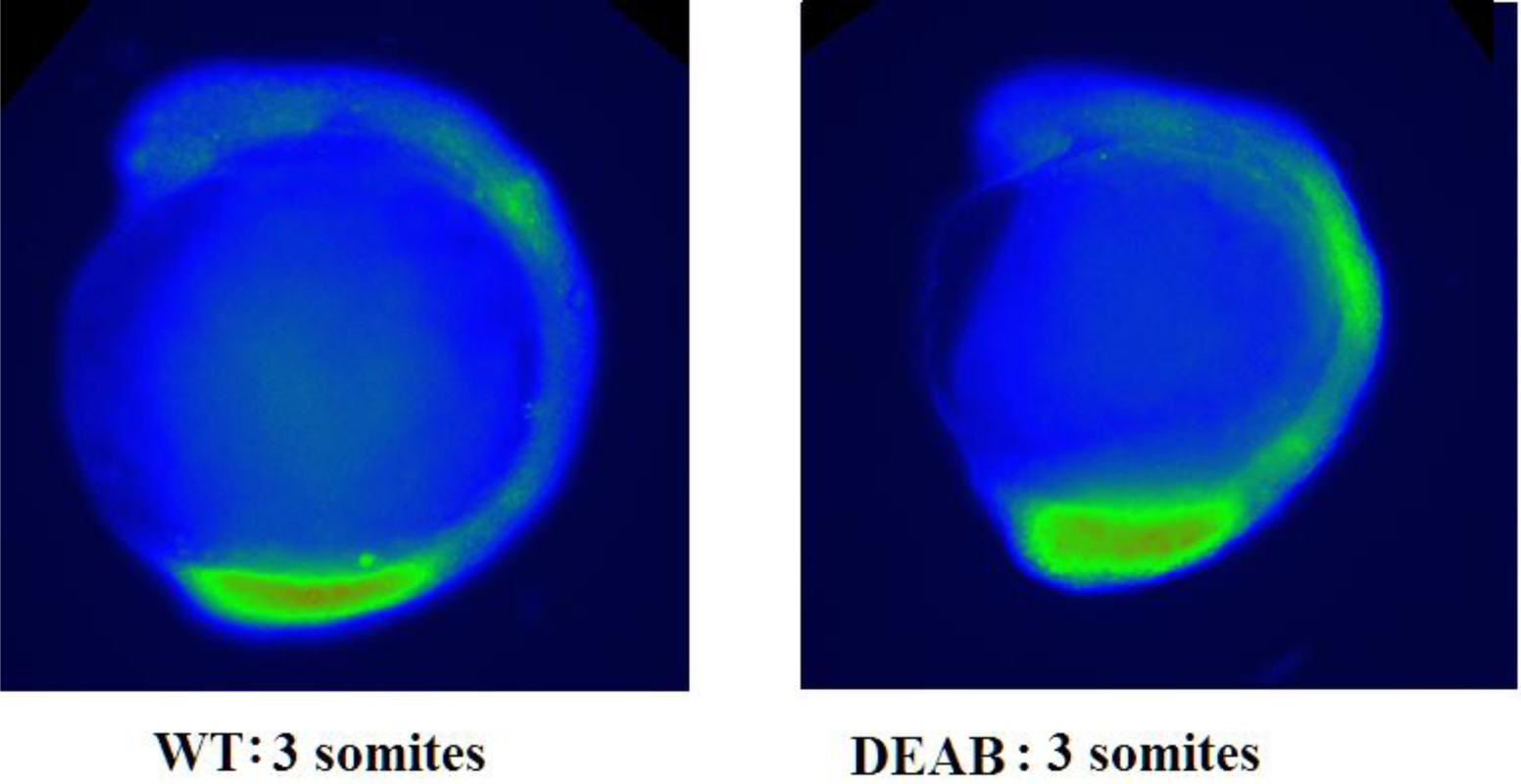
MapK domain of activity at 3 somites in WT and DEAB treated embryos. As predicted the domain of activity is larger in DEAB treated embryos.

**Fig. S3:**
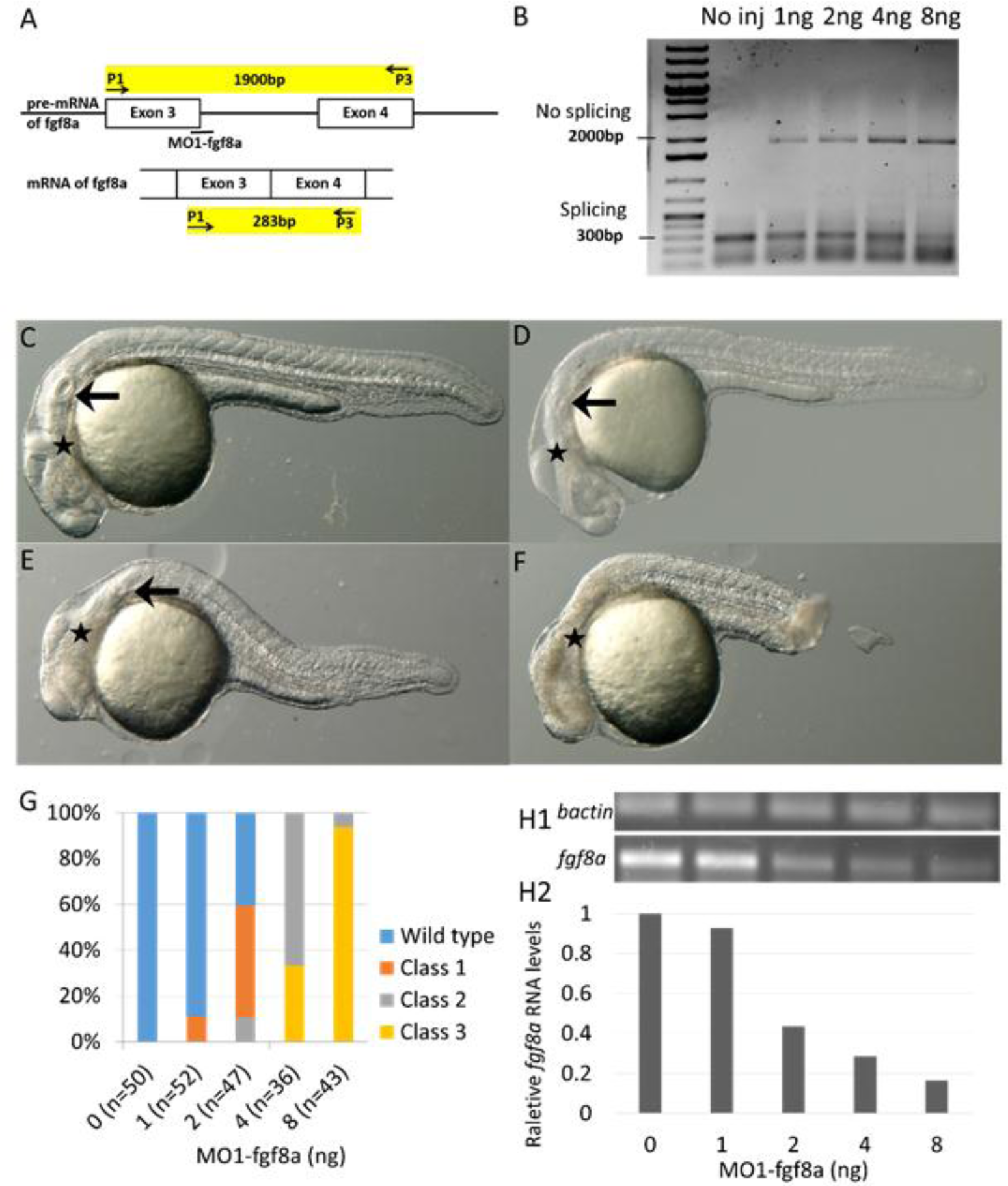
Phenotype of embryos injected with a morpholino (MO1-Fgf8) against the exon3-intron3 of Fgf8 (E3I3 in (Draper et al., 2001)) (A) and cDNA evidence of their dose-related effect (B): increase of the unspliced variant. (C-F): Lateral view of live embryo phenotypes at 24hpf reproduce those reported in (Draper et al., 2001): (C) Wild type embryo without injection show MHB (star) and otic vesicle (arrowhead); (D) Class I embryos exhibit clear loss of MHB and smaller otic vesicle; (E) Class II embryos show loss of MHB, smaller otic vesicles and somite defects; (F) Class III embryos present extensive tissue necrosis. (G) Phenotypic Class scoring at 24hpf in non-injected or 1ng, 2ng, 4ng, 8ng MO1-Fgf8 injected embryos. (H) Comparison of Fgf8 mRNA level in wild type and embryos injected with different amounts of MO1-Fgf8. (H) RT-PCR of fgf8 (H1) and β-actin (H2) Relative amount of Fgf8 mRNA compared to wild-type amount of Fgf8 mRNA transcripts.

**Fig. S4:**
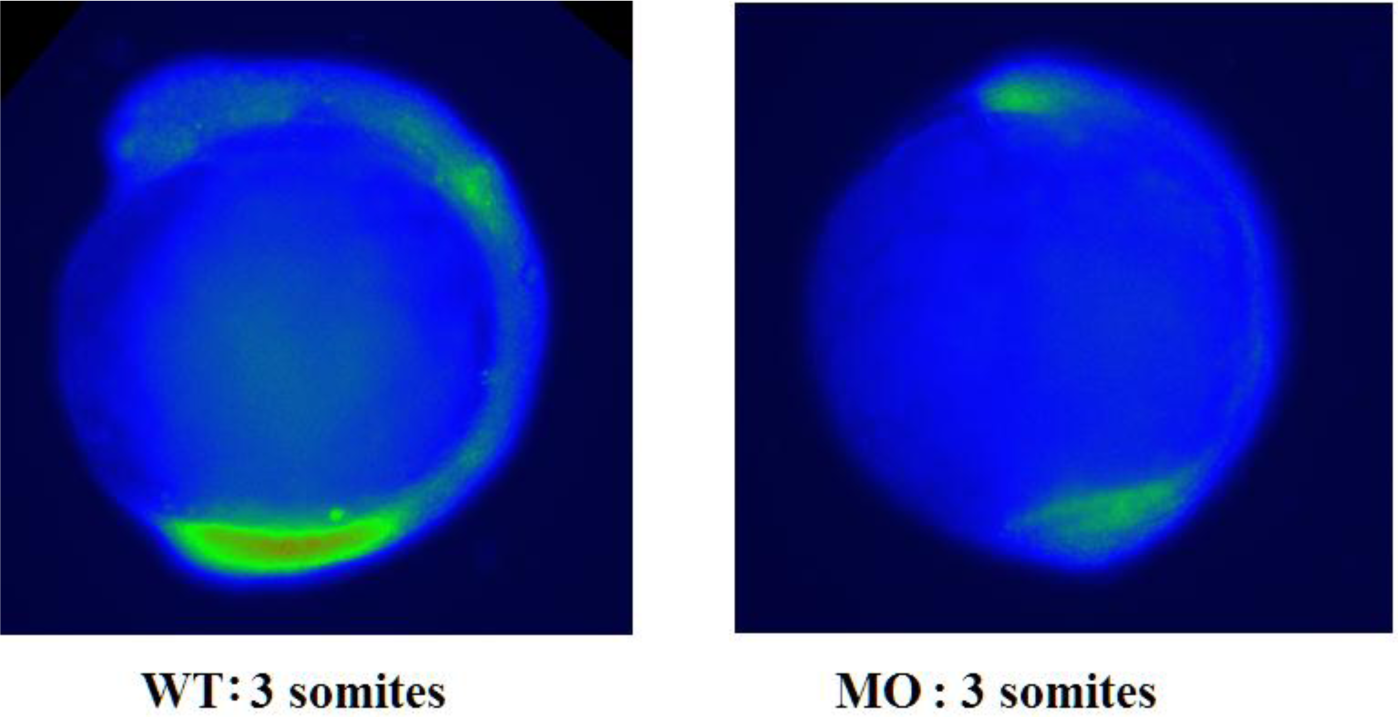
MapK domain of activity at 3 somites in WT and MO1-Fgf8 injected embryos. As predicted the domain of activity is smaller in MO1-Fgf8 injected embryos.

**Fig. S5:**
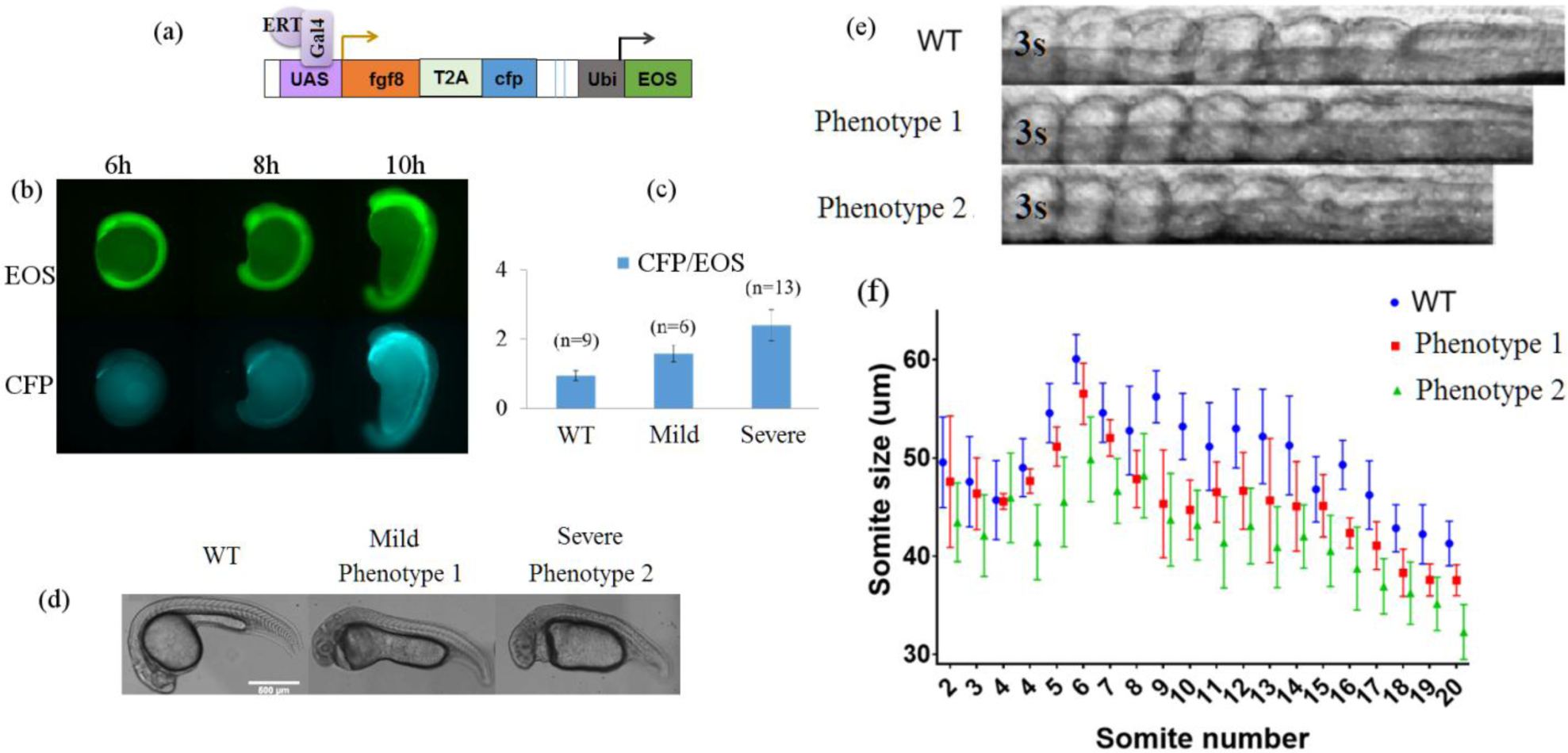
Phenotype observed upon induction of exogenous Fgf8. (a) the transgene uas:fgf8-T2A-cfp is turned on when a transcription factor Gal4-ERT is released from its cytoplasmic chaperone complex by binding of cyclofen (possibly photo-activated from caged cyclofen (cCyc) by UV light). A ubiquitous mEosFP informs on the presence of the transgene. (b) Embryos activated by 5 min UV (375nm) illumination of 6μM cCyc at 70% epiboly display an increase with time of CFP (and thus Fgf8) expression. (c) The ratio of CFP to mEosFP fluorescence (which is proportional to the level of Fgf8) correlates with the severity of developmental malformations which can be grouped at 24hpf in two classes (d): a mild phenotype (phenotype 1) characterized by enlarged heart and yolk and abnormal development of the head, and a severe phenotype (phenotype 2) characterized by enlarged heart, abnormal development of the head (with often no eyes), lost yolk extension and disordered late somites (>20s). (e) comparison of the initial stages of somitogenesis (3-8 somites) in WT and phenotypes 1 and 2: notice the smaller somites in the Fgf8 overexpressing embryos. (f) Somite size at onset in WT and phenotypes 1 and 2 as a function of the stage of somitogenesis. Notice the difference between early stages (before somite 6).characterized by an increase in the somite size and somitogenesis after 7 somites characterized by a slow decrease of the somite size (f).

**Fig. S6:**
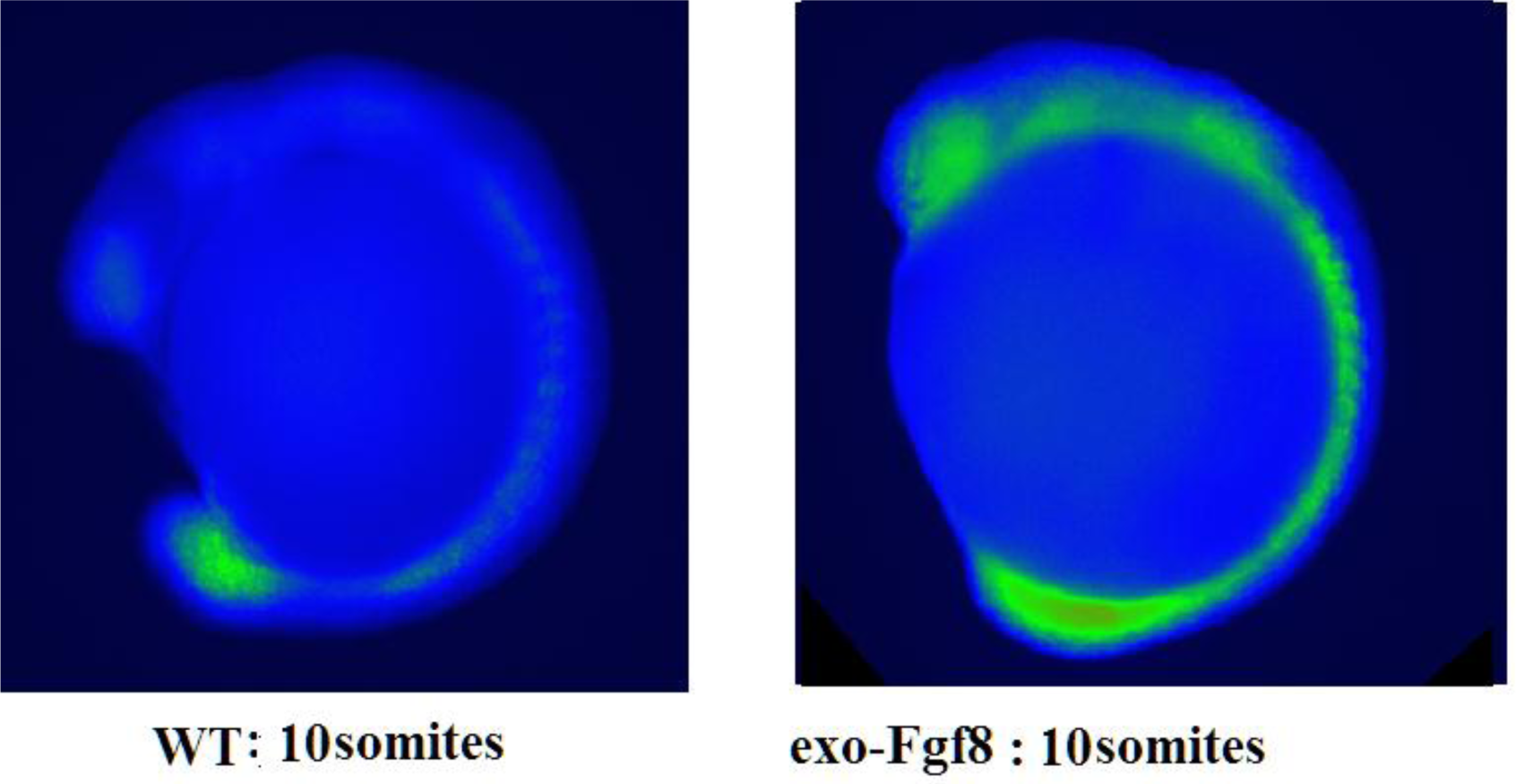
MapK domain of activity at 10 somites in WT and embryos with an exogenous source of Fgf8 (under control of a UAS promotor and a Gal-ERT transcription factor activated by cyclofen). As predicted the domain of activity is larger in embryos with an exogenous source of Fgf8.

**Fig. S7:**
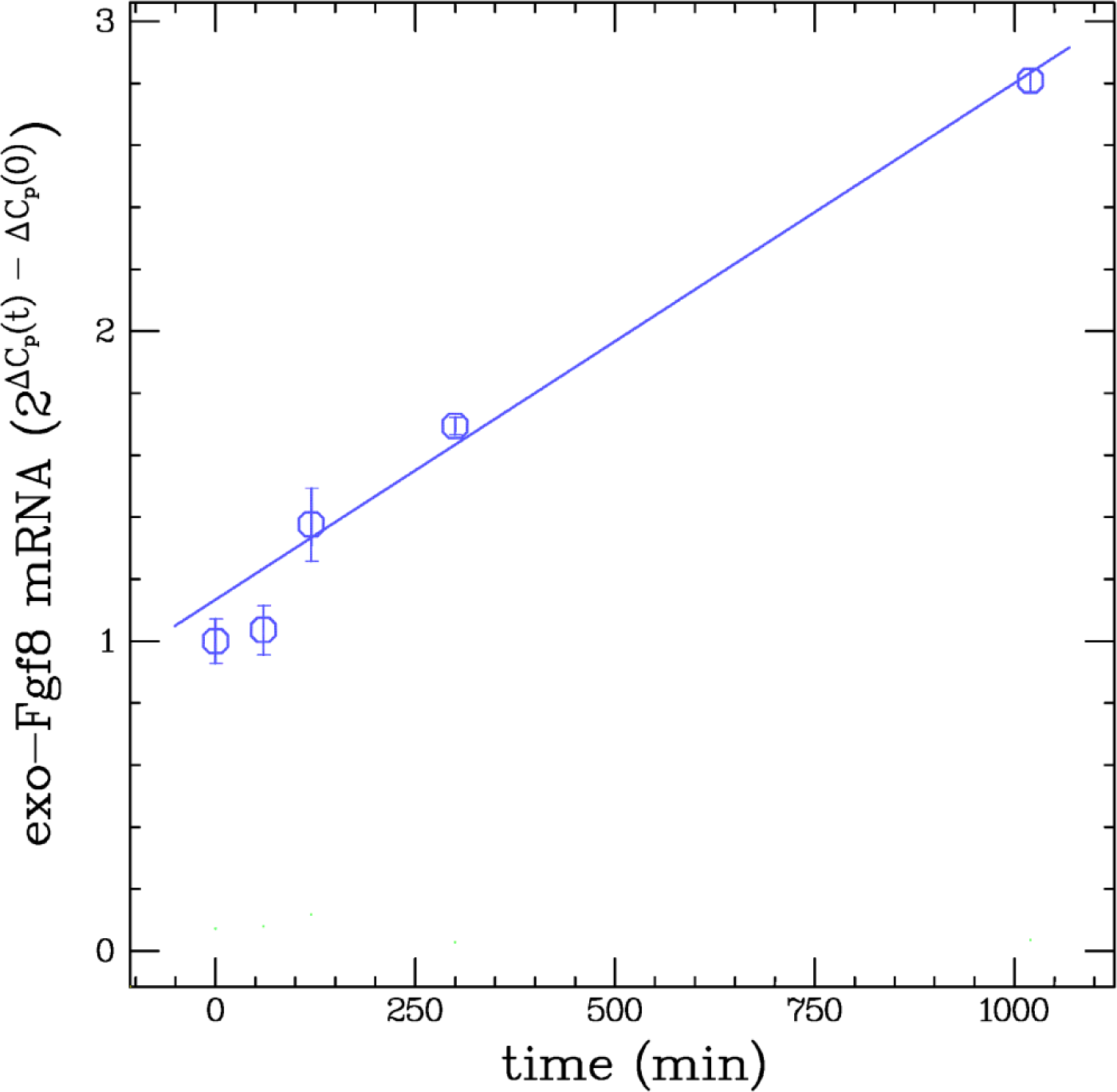
Increase of total Fgf8 mRNA in presence of an exogenous source of Fgf8 (see Fig.S5 above). At time t=0 (bud stage) the concentration of Fgf8 is contributed by the endogenous one. As time increases the endogenous concentration decreases rapidly (see Fig.4(a)) while the exogenous concentration increases with time. It typically doubles the initial endogenous concentration in 500 min (about 20 somite stage).

**Fig. S8:**
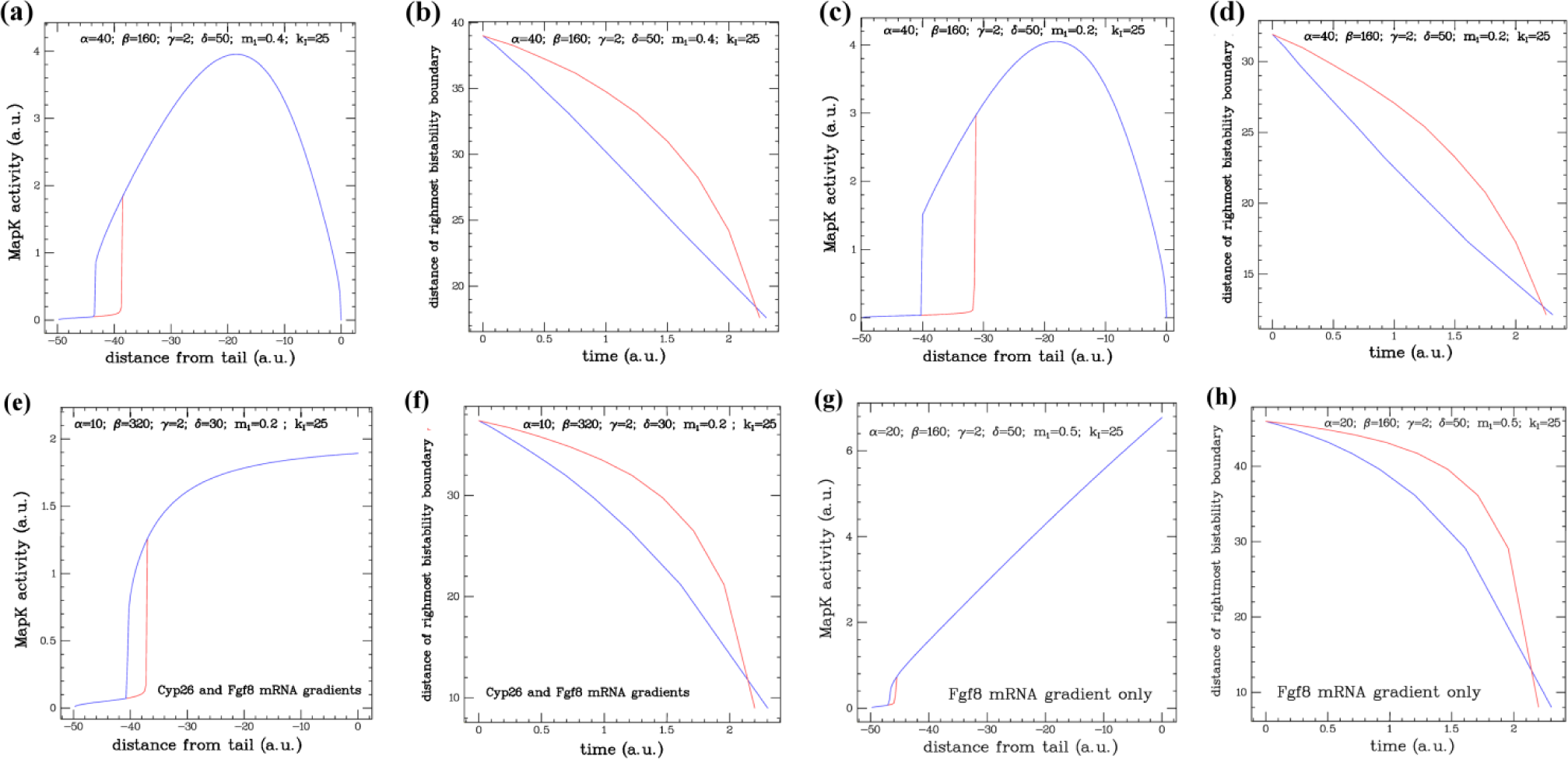
(a,c) MapK activity at steady-state from a simulation of mG_2_P model with the parameters shown in the figures and linear gradients of RaldH (parameter α) and Fgf8 mRNA (parameter γ). Notice the existence of a bistability window (between distances: −40 and −32 in Fig.2(c)). (b,d) Variation with time of the distance from the tail end (at 0) of the rightmost bistability boundary (red vertical line in (a,c)): assuming linear decrease with time of Fgf8 mRNA (red curve; t ∼ 1 - mF0(t)/mF(0)) or exponential decrease with time of Fgf8 mRNA (blue curve, mF0(t) = mF0(0) e^-t^). (e,f) Results of simulations with linear gradients of Cyp26 (parameter β) and Fgf8 mRNA (parameter γ). Notice that the MapK activity levels off at the tail end though a bistability window is still present. (g,h) Results of simulations with a linear gradient of Fgf8 mRNA (parameter γ) only. Notice the increasing MapK activity level and the increased PSM shrinkage rate at the tail end (even assuming an exponential decay with time of FgF8 mRNA).

**Fig. S9:**
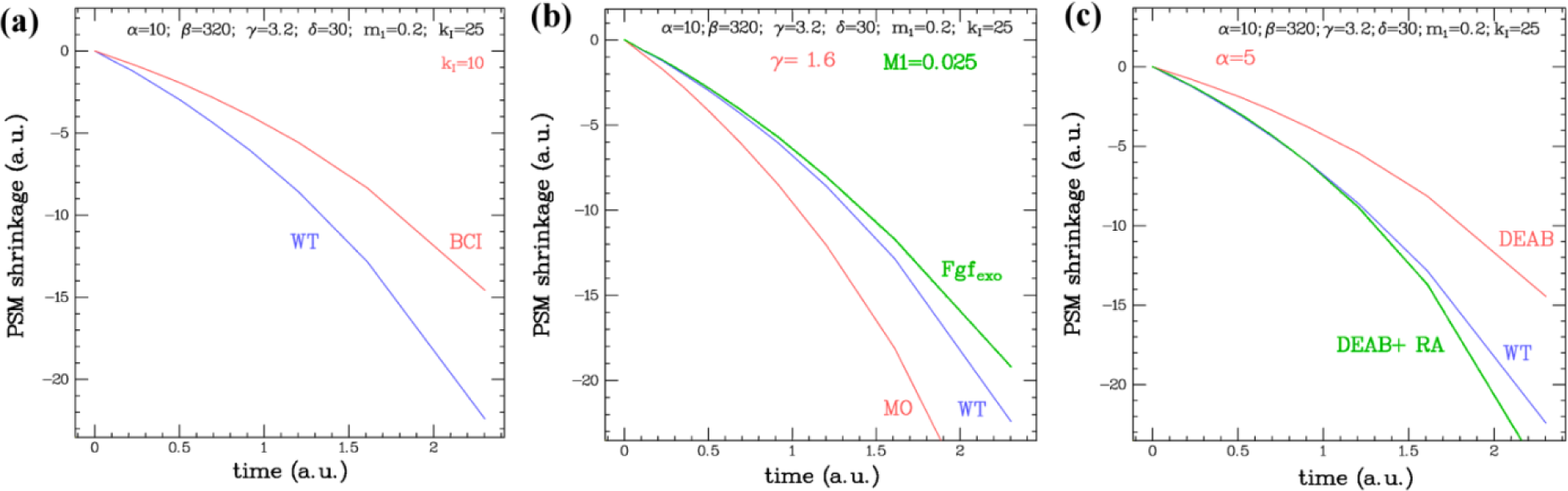
Simulation of the mG2P model with linear gradients of Cyp26 (parameter β) and Fgf8 mRNA (parameter γ) Results labeled WT are the results of a simulation with the parameters shown at the top. (a) Results simulating an inhibition of Mkp3 by BCI (decrease of parameter *k*_*I*_ by a factor 2.5). The simulation is in qualitative agreement with the observations (Fig.5).(b) Results simulating the effects of Morpholinos against Fgf8 (decrease of parameter γ by a factor 2) and the presence of an increasing concentration of exogenous Fgf8 mRNA. The results are in qualitative agreement with the observations (Fig.7), even though the MO effect is overstated. (c) Results simulating an inhibition of RaldH by DEAB (decrease in parameter α by a factor 2). These results are qualitatively different from the observations (see Fig.6): in presence of DEAB the PSM rate of shrinkage is decreased in contradiction with the observations.

## Supplementary experimental data

1).RA positively regulate Fgf8.

2).MapK domain of activity in presence of DEAB

3).MO1-Fgf8 effect on phenotype

4).MapK domain of activity in presence of MO1-Fgf8

5).Effect of exogenous Fgf8 expression on developmental phenotype.

6).MapK domain of activity in presence of an exogenous source of Fgf8

7).qPCR of exogenous Fgf8

## The mG_2_P model

The modified G_2_P model proposed here is shown below.

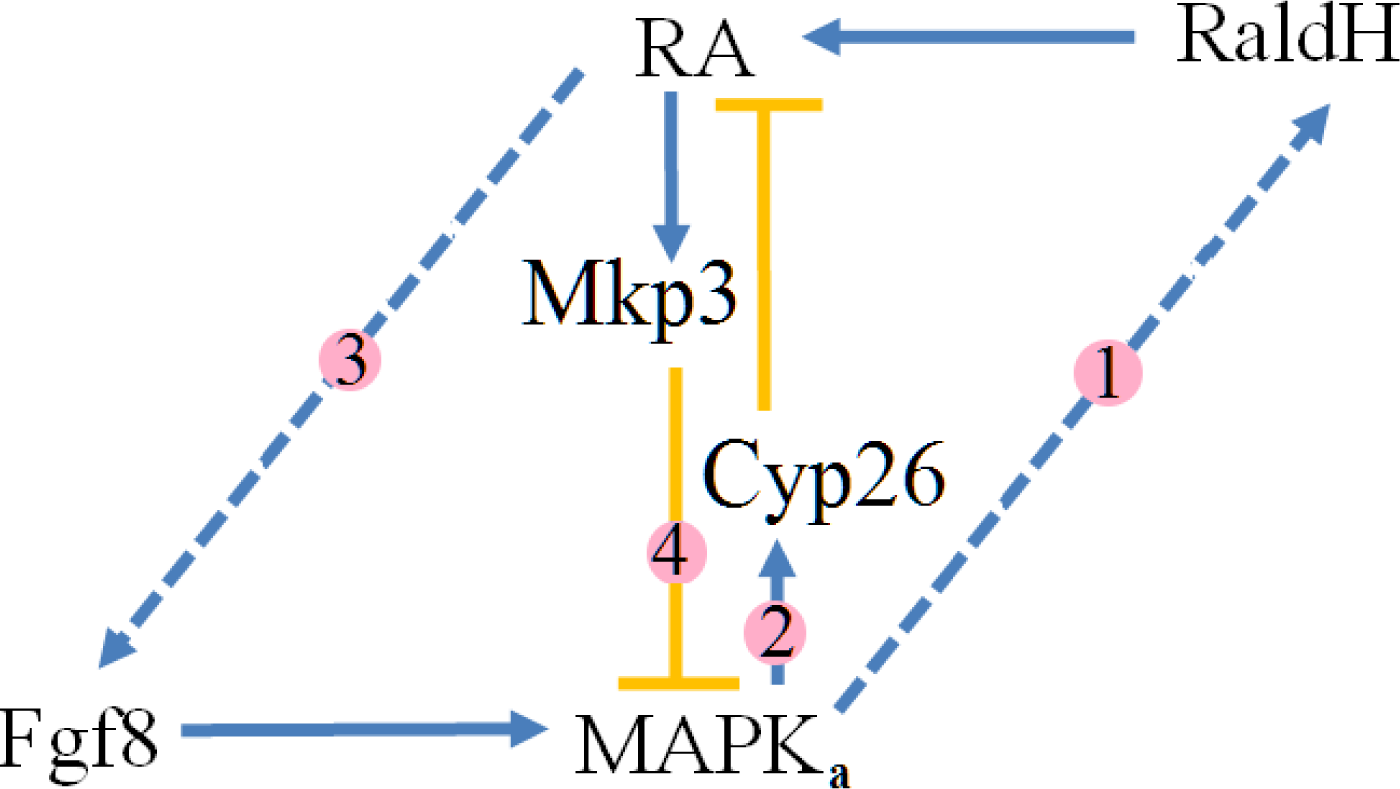

In the moving frame of the tail, the equations describing that molecular network are as follows.

Retinoic acid *[RA]* is synthesized from retinal by Retinal dehydrogenase (RaldH) at rate *v*_*sr*_ and degraded at rate *k*_*dr*_ by Cyp26 (at conc. *C*) and at a basal rate measured by the kinetic constant *k*_*d1*_.

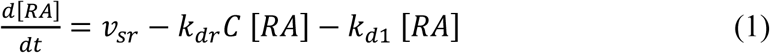

The reported positive feedback of the active (phosphorylated) form of ***MapK*** (at concentration ***MK*_*a*_**) on RaldH (see pathway 1 above; Hamada et al.,2006; Shimozono et al., 2013) is implemented as a Hill equation with half occupation concentration *Mk*_*r*_ and a non-cooperative binding (Hill coefficient = 1):

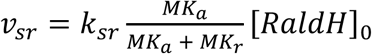

The mRNA of Cyp26 (*mC*) is under positive feedback of ***MK*_*a*_** (see pathway 2 above) and is degraded at rate *k*_*d2*_

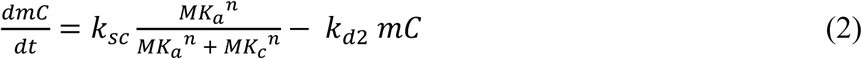

Where we assumed following (Goldbeter et al., 2007) a non-linear (*n=2*) Hill equation. This non-linearity is essential for bi-stability.

Cyp26 is synthesized from its mRNA at rate *k*_*s3*_ and degraded at rate *k*_*d3*_.

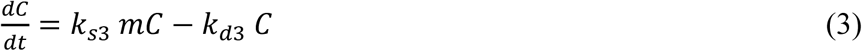

Fgf8 is synthesized from its mRNA (*mF*) at a rate *k*_*s4*_ and degraded at rate *k*_*d4*_

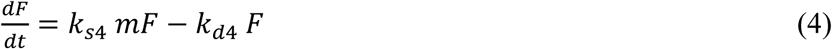

To implement the observed positive feedback of [RA] on *mF* (see pathway 3 above) we use a Hill equation with non-cooperative binding of [RA] and half occupation *R*_*0*_.

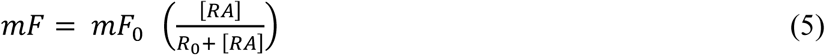

The core of the network is the ***MapK*** which is activated (phosphorylated) by the ***Fgf8*** pathway and inhibited by ***[RA]*** via its action on Mkp3 (see pathway 4 above):

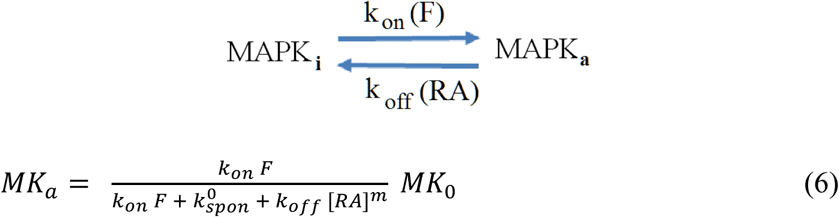

Where *MK*_*0*_ is the total MapK concentration, ***k*_*on*_ *F*** is the Fgf8 dependent rate of MapK phosphorylation and ***k*^*0*^_*spon*_** and ***k*_*off*_ *[RA]*^*m*^** are respectively the RA independent and dependent rates of MapK dephosphorylation. A non-linear Hill coefficient (*m=2*) is required for bi-stability.

These equations were simulated (using MATLAB) and at steady state they can reach one or two stable states depending on the initial conditions and the various parameters. Actually at steady-state the above equations simplify enormously. Defining:

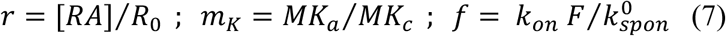

yields:

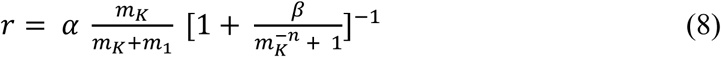

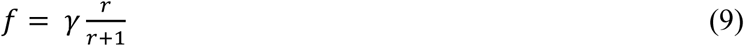

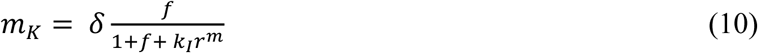

With:

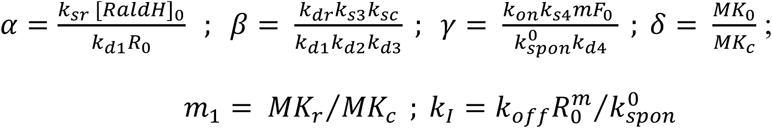

Thus at steady state instead of 18 parameters the system depends on only 6 non-dimensional ones (as can be verified by comparing the results of simulations of Eqs.(1-6) with the steady-state predictions Eqs.(8-10)). In the limit 1≪ r; *m*1≪ *m*_*k*_ (negligible pathways 1 and 3), the above model is formally identical to the G_2_P model (with MapK playing the role of Fgf8).

If we let some of the parameters in the above model depend on the distance from the tail end, then there might exist along the PSM a window of bistability where two stable solutions are possible with a high or a low MapK activity (variable *m*_*K*_ in Eq.(10)). In the comparison with the experimental data we have assumed following (20) that RalDH and the Fgf8 mRNA displayed a monotonous gradient, i.e.:

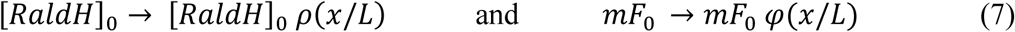

Where 0 < *ρ (u), φ(u)* < 1 and *L* is a typical scale assumed to be constant during somitogenesis. For simplicity we have set: *φ(u)* = *u* and *ρ (u), =* 1 - *φ(u)* We have simulated the mG_2_P model with exponential gradients in Fgf8 and RaldH, i.e *φ(u)* = 1 – *e* ^*–u /u*^_*0*_ or with *ρ(u) =* 1 (i.e. in presence of a Fgf8 mRNA gradient only) or in presence of a Cyp26 mRNA gradient (*k*_*sc*_ → *k*_*sc*_ *φ (x/L*)), instead of a RaldH gradient, see Fig.S8. While a bi-stability window can always be found, the PSM rate of shrinkage is not constant in these different situations (see below).

When inducing the expression of exogenous Fgf8 we add to *mF*_*0*_ an extra term *mF*_*exo*_ *(t),* which varies on a much longer time-scale than the relaxation of the system to its steady-state value (adiabatic approximation). Similarly in presence of an external concentration of RA, we add a constant term *RA*_*ext*_ to *[RA].* A typical set of parameters (similar to the ones mentioned in (Goldbeter et al., 2007)) is shown in the table below:

The model was run on MATLAB with the parameters shown in table I (and others) and at steady state it clearly displays a bistability window (in the range −40 < x < −32 in Fig.S8(c)), see Fig.S8(a,c) below.

**Table 1:** Typical set of parameters used in the simulation of model mG_2_P.

|  |  |  |  |  |  |
| --- | --- | --- | --- | --- | --- |
| $k_{sr} = 1 \text{ min}^{-1}$ | $MK_r = 0.4 \text{ nM}$ | $MK_c = 1 \text{ nM}$ | $MK_0 = 50 \text{ nM}$ | $[RaldH]_0 = 20 \text{ nM}$ | $R_0 = 10 \text{ nM}$ |
| $k_{dr} = 0.1 \text{ nM}^{-1} \text{ min}^{-1}$ | $k_{sc} = 2 \text{ nM min}^{-1}$ | $k_{s3} = 0.12 \text{ min}^{-1}$ | $k_{s4} = 0.1 \text{ min}^{-1}$ | $k_{on} = 1 \text{ nM}^{-1} \text{ min}^{-1}$ | $mF_0 = 40 \text{ nM}$ |
| $k_{d1} = 0.05 \text{ min}^{-1}$ | $k_{d2} = 0.05 \text{ min}^{-1}$ | $k_{d3} = 0.06 \text{ min}^{-1}$ | $k_{d4} = 1 \text{ min}^{-1}$ | $k_{spon} = 2 \text{ min}^{-1}$ | $k_{off} = 0.5 \text{ nM}^{-2} \text{ min}^{-1}$ |
| $\alpha = 40$ | $\beta = 160$ | $\gamma = 2$ | $\delta = 50$ | $m_1 = 0.4$ | $k_I = 25$ |

According to this model, at fixed times (i.e. a set phase of the somitogenetic clock) the undifferentiated posterior PSM (high MapK activity state, blue curve in Fig.S8(a,c,e,f)) is switched in its bistability window into a somite committed state (low MapK activity state, red curve in Fig.S8(a,c,e,f)). Within the present model this can easily be accounted for by a wave of MapK inhibition (for example by Sprouty, Hayashi et al., 2009) reaching the bistability window and switching MapK to its low bistability state. The MapK activity boundary then jumps to its rightmost stability boundary. These coordinated jumps of the MapK activity domain have been reported recently (Sari et al., 2018).

To address the observation of PSM shrinkage during somitogenesis, we used the above model to see how the PSM would shrink with time if the concentration of Fgf8 mRNA (parameter *mF*_*0*_ above) decreased with time (on a timescale longer than the time to reach steady state, i.e. in the adiabatic approximation). As can be seen in Fig.S8(b,d) an almost linear decrease (constant rate of shrinkage) is observed if the concentration decreases exponentially with time, but not if it decreases linearly with time (in that case the rate of shrinkage increases with time). This is a quite generic feature of the model (it does not depend much on a variation of the model parameters, compare Fig.S8(b), Fig.S8(d) and Fig.S8(f))

Next, we investigated how the model predictions vary in presence of definite perturbations such as:

1. The presence of BCI, an inhibitor of Mkp3 that effectively decreases *k*_*off*_ (dimensionless parameter *k*_*I*_).
2. DEAB, an inhibitor of RaldH that affects the rate of RA synthesis *k*_*sr*_ (dimensionless parameter α) with or without an external source of RA, that can be incorporated easily in the simulations.
3. A morpholino against Fgf8 which effectively affects his concentration of Fgf8 mRNA *mF*_*0*_ (dimensionless parameter γ).
4. The presence of an exogenous source of Fgf8 that can be turned on at the beginning of somitogenesis. That experiment is modeled by adding a time dependent source of Fgf8 mRNA in eq.(7) above: Fgf_ext_ = M1*t (in the simulations presented here M1=0.005 *mF*_*0*_ per unit time).

The results are presented in the text. The spatial scale of the simulations was set by matching the size of the region with high MapK activity (on the right of the leftmost bistability boundary in Fig.S8(a,c)) to the experimentally measured size of the PSM in unperturbed embryos at 7 somites (which defines time t=0). The temporal scale was set by rescaling the time scale of Fgf8 mRNA decrease in the simulation (τ_sim_ = 1) to the time-scale of the same decrease in the experiment (estimated from qPCR measurement of Fgf8 mRNA concentration to be: τ_fgf8_ ≅ 8 somitic periods), see Fig.4. The results with linear gradients of RaldH (parameter α) and Fgf8 mRNA (parameter γ) are shown in the main text where they are compared to our observations.

We have also investigated other possible gradients, such as a gradient of Fgf8 mRNA only, or a gradient of Fgf8 mRNA and Cyp26 transcription rate (*k*_*sc*_ or parameter β). These cases seem to be incompatible with some of our observations. For example a single linear gradient of Fgf8 mRNA, while compatible with the existence of a bi-stability window (Fig.S8(g)) and thus differentiation into somites, is incompatible with an almost constant rate of shrinkage of the PSM (assuming a Fgf8 mRNA exponentially decaying over an 8 somite time scale), see Fig. S8(h). It also yields a linear gradient of MapK activity (Fig.S8(g)) which is not observed.

Results with dual co-linear gradients of Fgf8 mRNA (parameter γ) and Cyp26 transcription rate (parameter β) are compatible with a bi-stability window, an almost constant rate of shrinkage of the PSM and our observations of slower PSM shrinkage rate in presence of BCI (an inhibitor of Mkp3), Fig.S9(a), or morpholinos against Fgf8 and a faster rate in presence of an increasing concentration of exogenous Fgf8 mRNA, Fig.S9(b). The results of this simulations are however incompatible with a faster shrinkage rate in presence of an inhibitor of RaldH (DEAB), compare Fig.S9(c) and Fig.6.

